# A Bayesian method to cluster single-cell RNA sequencing data using Copy Number Alterations

**DOI:** 10.1101/2021.02.02.429335

**Authors:** Salvatore Milite, Riccardo Bergamin, Lucrezia Patruno, Nicola Calonaci, Giulio Caravagna

## Abstract

**Motivation:** Cancers are composed by several heterogeneous subpopulations, each one harbouring different genetic and epigenetic somatic alterations that contribute to disease onset and therapy response. In recent years, copy number alterations leading to tumour aneuploidy have been identified as potential key drivers of such populations, but the definition of the precise makeup of cancer subclones from sequencing assays remains challenging. In the end, little is known about the mapping between complex copy number alterations and their effect on cancer phenotypes.

**Results:** We introduce CONGAS, a Bayesian probabilistic method to phase bulk DNA and single-cell RNA measurements from independent assays. CONGAS jointly identifies clusters of single cells with subclonal copy number alterations, and differences in RNA expression. The model builds statistical priors leveraging bulk DNA sequencing data, does not require a normal reference and scales fast thanks to a GPU backend and variational inference. We test CONGAS on both simulated and real data, and find that it can determine the tumour subclonal composition at the single-cell level together with clone-specific RNA phenotypes in tumour data generated from both 10x and Smart-Seq assays.

**Availability:** CONGAS is available as 2 packages: CONGAS (https://github.com/caravagnalab/congas), which implements the model in Python, and RCONGAS (https://caravagnalab.github.io/rcongas/), which provides R functions to process inputs, outputs, and run CONGAS fits. The analysis of real data and scripts to generate figures of this paper are available via RCONGAS; code associated to simulations is available at https://github.com/caravagnalab/rcongas_test.

**Supplementary information:** Supplementary data are available at *Bioinformatics* online.

## 1 Introduction

Cancers grow from a single cell, in an evolutionary process modulated by selective forces that act upon cancer genotypes and phenotypes (Greaves and Maley, 2012; McGranahan and Swanton, 2015). The fuel to cancer evolution is genotypic and phenotypic cellular heterogeneity, and much is yet to be understood regarding its effect on evolution and response to therapy (McGranahan and Swanton, 2017; Turajlic *et al*., 2019). Notably, the heterogeneity observed in cancer can also be produced during normal tissue development, and therefore the quest for understanding heterogeneity has implications far beyond cancer (Martincorena *et al*., 2015; Martincorena, 2019).

While the evolutionary principle of cancer growth is intuitive to conceptualise and replicate in-vivo (Acar *et al*., 2020), it is still hard to precisely measure clonal evolution using sequencing technologies (Caravagna, 2020). Even if popular single-omic assays from 10x and Smart-Seq achieve higher resolution than bulk counterparts (Picelli *et al*., 2014; Wang *et al*., 2019), their analysis poses many challenges (Lähnemann *et al*., 2020). Nowadays, much hope is put into multi-omics technologies that probe multiple molecules from the same cell (Macaulay *et al*., 2015). Multi-omics data explicitly gather DNA and RNA measurements per cell; unfortunately, however, such assays are still too expensive to scale to more than hundreds of cells. An interesting opportunity is attempting the integration of different types of single-omic assays that, individually, already scale to thousands of cells. At least conceptually, the statistical integration of independent assays comes from mapping one dataset on top of another, leveraging a quantitative model for the relation between the sequenced molecules (e.g., we may wish to predict DNA from RNA, or vice versa).

In this work we develop a Bayesian method for total Copy Number Genotyping from single-cells (CONGAS), which integrates total Copy Number Alterations (CNAs) obtained from bulk DNA sequencing and single-cell RNA data (scRNA) from independent cells (Figure 1a). Our method is similar to clonealign, which uses two single-cell assays to assign scRNA profiles to tumour clones predetermined from low-pass single-cell DNA sequencing (Campbell *et al*., 2019). CONGAS and clonealign conceptualise the same linear model to link total CNAs - i.e., the sum of the major and minor allele copies (Househam *et al*., 2021) - with RNA counts, but while clonealign fixes the set of clones from its input and is therefore supervised (Supplementary Figure S1), CONGAS is unsupervised and finds new clusters by leveraging Bayesian priors from the input bulk (Figure 1b). Precisely, CONGAS uses input CNAs to define the genome segmentation and parametrise a prior for total CNAs of each segment - then each cluster has its posterior distribution over the ploidy of the segments. We note that the extra input for CONGAS can be generated from a routine low-pass bulk DNA assay, which is much cheaper then the single-cell counterpart required by clonealign.

**Figure 1.**
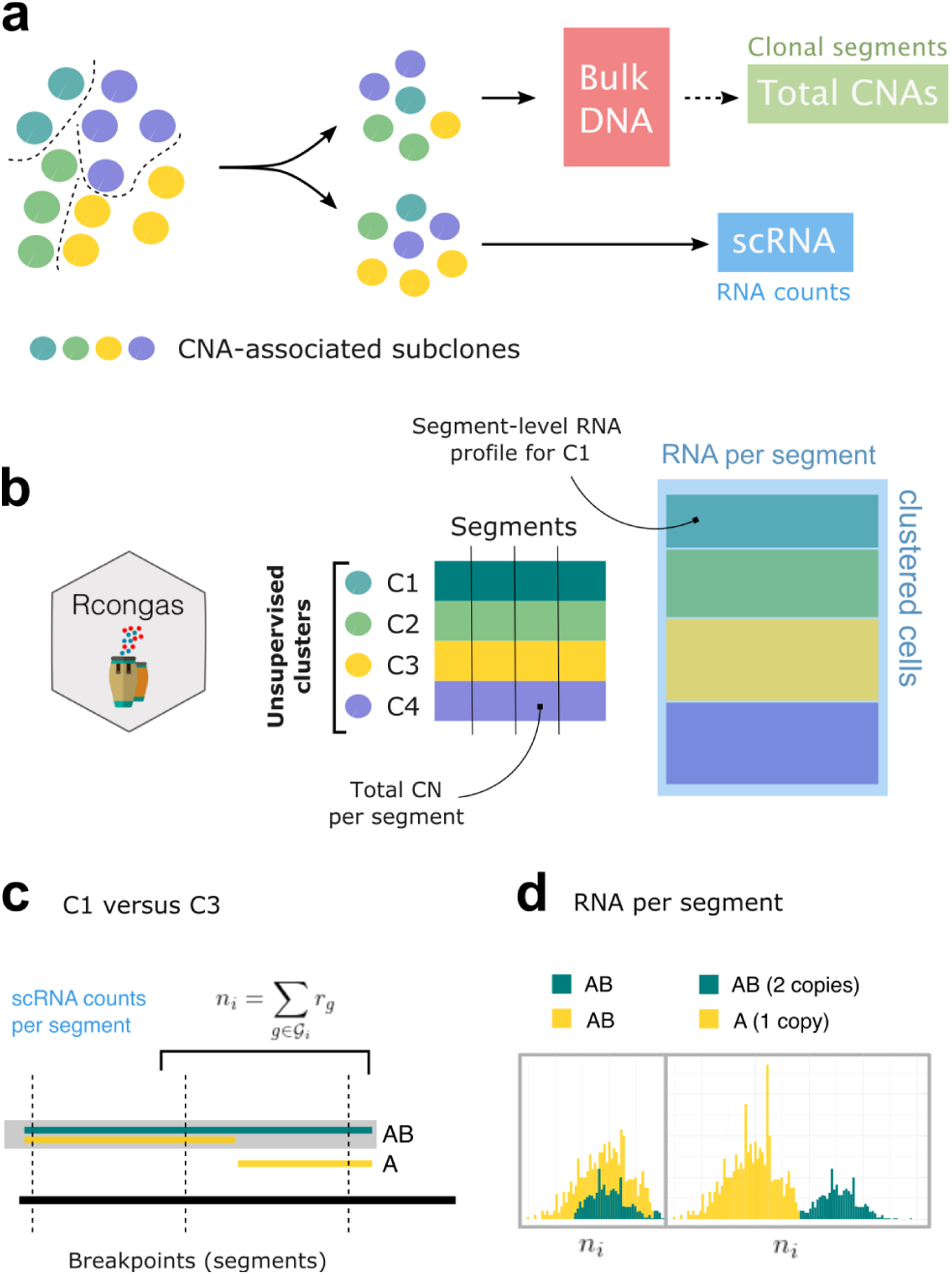
**a**. CONGAS works with 1) total CNA data (ploidy values per segment) from a bulk DNA assay and 2) single-cell RNA sequencing data (scRNA). The two assays are generated from independent cells of the same starting sample. The aim is to identify CNA-associated subclones from RNA counts. **b**. CONGAS is a Bayesian unsupervised method to identify clusters of cells whose differences in scRNA counts can be explained by total CNAs. Subclonal CNAs are here inferred at the resolution of the input segments. **c**,**d**. Assume subclones C1 and C3 differ for a portion of DNA (right segment): C3 has a subclonal LOH (A genotype), where C2 is heterozygous diploid (AB genotype). CONGAS identifies CNAs by examining total RNA counts mapped to segments: subclone C3 shows fewer RNA counts on the deleted segment, and the subclones have similar RNA counts on the segment where both clones are heterozygous diploid (left segment).

With CONGAS we formulate an unsupervised clustering problem: we seek to group cells with segment-level RNA profiles that can be explained by similar CNAs (Figure 1c,d), inferring CNAs and clusters jointly. There are methods that are alternative to CONGAS, for instance InferCNV, HoneyBADGER, CASPER and copyKAT, that detect CNAs by segmenting scRNA counts (Fan *et al*., 2018; Serin Harmanci *et al*., 2020). These methods however decouple CNA detection from clustering, requiring to select the number of optimal clusters with some heuristic. Instead, CONGAS detects subclonal CNAs and clusters cells in a unified model, therefore integrating uncertainty with its Bayesian formulation. Compared to some of the alternative methods CONGAS also has the advantage of working without reference scRNA expression; this avoids using RNA tissue databases, or requiring normal cells in the input scRNA assay. Therefore CONGAS can be applicable in designs where the normal signal is difficult to obtain, e.g., with cancer organoids. CONGAS can associate CNA-associated cancer subclones to transcriptomic profiles, providing an explicit mapping between genotype and phenotype at the clone level. This is particularly important in cancer, where we want to characterise how chromosomal instability drives tumour evolution (Watkins *et al*., 2020), or, where we want to understand how pre-cancerous cells can be causally linked to the onset of cancer (Martincorena, 2019).

## 2 Methods

The aim of CONGAS is to statistically integrate DNA and RNA measures for every cell, deriving a measure of total DNA abundance per segment (i.e., total copy number) and RNA counts per cell. This accounts for emulating a DNA-RNA multi-omics assay, which we use to cluster cells whose RNA profile can be explained by similar copy numbers.

### 2.1 CONGAS

CONGA is a Bayesian method that “genotypes” bulk CNAs on top of scRNA data; The term genotyping elicits the use in CONGAS of an input set of CNAs obtained from bulk DNA sequencing, here used to create Bayesian priors. A vector of input total copy number profiles drives the calling of subclonal CNAs from single-cells, in a way that new CNAs can be obtained as ploidy changes with fixed breakpoints. In particular, breakpoints are used to pool RNA counts per segment, and bulk-level total copy numbers constitute a Bayesian prior per segment. Therefore, the model is able to infer variations of single-cell CNAs around the input bulk. The CONGAS likelihood is a mixture of *K* > 0 Poisson distributions for scRNA counts per-segment, and works also with data normalised in common units; the likelihood is conditioned on the latent CNAs that we infer for each of the *K* cluster, and normalises counts for library size and number of genes per segment if required.

A low-pass bulk DNA assay to generate the input CNAs required for CONGAS is inexpensive. If this is unavailable, RNA segmentation can also be attempted, or an arm-level segmentation with constant ploidy 2 can be used to detect large CNAs. Input CNAs simplify the statistical inference problem and avoid the segmentation of noisy single-cell RNA data. The model chases subclonal populations that show different total CNAs at the resolution of the input segments. For instance, it can detect a subclonal population underlying a loss of heterozygosity (Figure 1b). After pooling RNA counts on every segment (Figure 1c), under a linear model that links DNA abundance to RNA counts (Campbell *et al*., 2019), we use Poisson distributions parameterized by unobserved copy number values to explain counts (Figure 1d). By this definition clonal CNAs - i.e., present in 100% of the input cells - show the same RNA signal and cannot be detected unless normal cells are in the sample (e.g., tumour versus normal). Nonetheless, the difference across subpopulations can be still captured wherever present (e.g., tumour subpopulation 1 versus tumour subpopulation 2).

The model likelihood with the usual independence assumption among cells and segments is

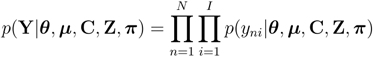

Here **Y** is the *N* × *I* input data matrix of RNA counts, which describe *N* sequenced cells and *I* input segments (mapped anywhere on the genome). Counts on a segment *y*_*n,i*_ are summed up by pooling all genes that map to the segment; with cumulative counts we rarely observe 0-counts segments, which allows us to avoid zero-inflated distributions (Sarkar and Stephens 2020). The segment likelihood is

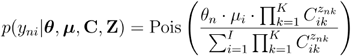

where the model uses θ_*n*_, a Gamma-distributed latent variable which models the library size for cell *n*, and μ_*i*_ for the number of genes in segment *i* (a constant determined from data). In CONGAS **C** is the clone CNA profile for *k* clones, where each clone is defined by *I* segments and associated CNAs; the prior for **C** is a log-transform of a normal distribution, consistently with the fact that ploidies are positive values. In this formulation **Z** are the *N* × *k* latent variables that assign cells to clusters, and π the *k*-dimensional mixing proportions (Supplementary Figure S2). Based on this modelling idea we also built alternative models that can process input data when these are already corrected for library size (e.g., in units of transcripts or read fragments per million) using Gaussian likelihoods; see Supplementary Materials.

Another way of thinking of the denominator in the formula is, given that all the effects are linear, as a matrix decomposition of the input. Note that here the denominator is omitted.

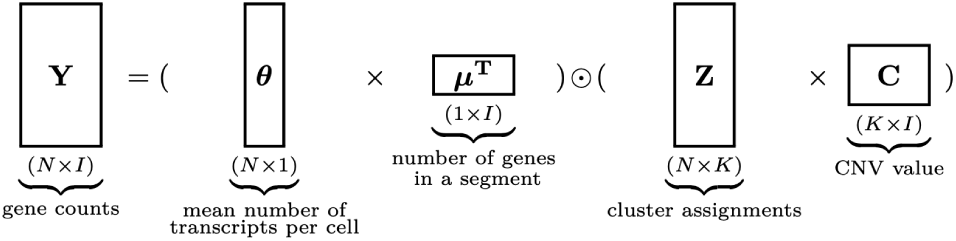

CONGAS parameters are learnt via stochastic variational inference (Blei *et al*., 2017). The model joint distribution can be factored as

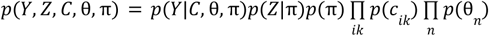

and in the variational framework latent variables are approximated as variational distributions *q*(*Z, C*, θ, π), supposed to be independent and factorizable. The prior distributions for our latent variables are:

- *p*(*c*_*ik*_)~*LogNorm*(*m*_*ik*_, *v*), where *m*_*ik*_ is the input CNA value from bulk DNA, and the variance *v* governs how far the actual CNAs can be compared to input (default *v* = 0. 5);
- *p*(θ_*n*_)~*Gamma*(*e*_*s*_, *e*_*r*_), a scarcely informative prior that works well in most cases (default *e*_*s*_ = 3, *e*_*r*_ = 1);
- *p*(π)~*Dirichlet*(*r*), a prior over cluster distributions, by default all assumed to have equal proportions (i.e. *r* = 1/*k*).

The CONGAS model is implemented in 2 open-source R/Python packages. One, called CONGAS, implements the model in the Pyro probabilistic programming language, a backend that allows running on both CPU and GPU (Bingham *et al*., 2019). A frontend R package, called RCONGAS, provides functions for data pre-processing, visualisation and model inference.

## 3 Results

### 3.1 Synthetic simulations

#### Generative model

We tested CONGAS by simulating synthetic data from its generative model, emulating a common 10x assay (1000 cells) for tumours of various complexity. Overall, we could retrieve the generative model in a number of scenarios for tumours with up to 5 CNA-associated subclones, evolving both linearly and branching (Figure 2a). The performance was measured via the Adjusted Rand Index (ARI), the ratio of agreements over disagreements in cell clustering assignments, and was consistent with other information-theoretic scores (Supplementary Figure S3). Clustering assignments were stable across a number of configurations of different subclonal complexity (Figure 2b).

**Figure 2.**
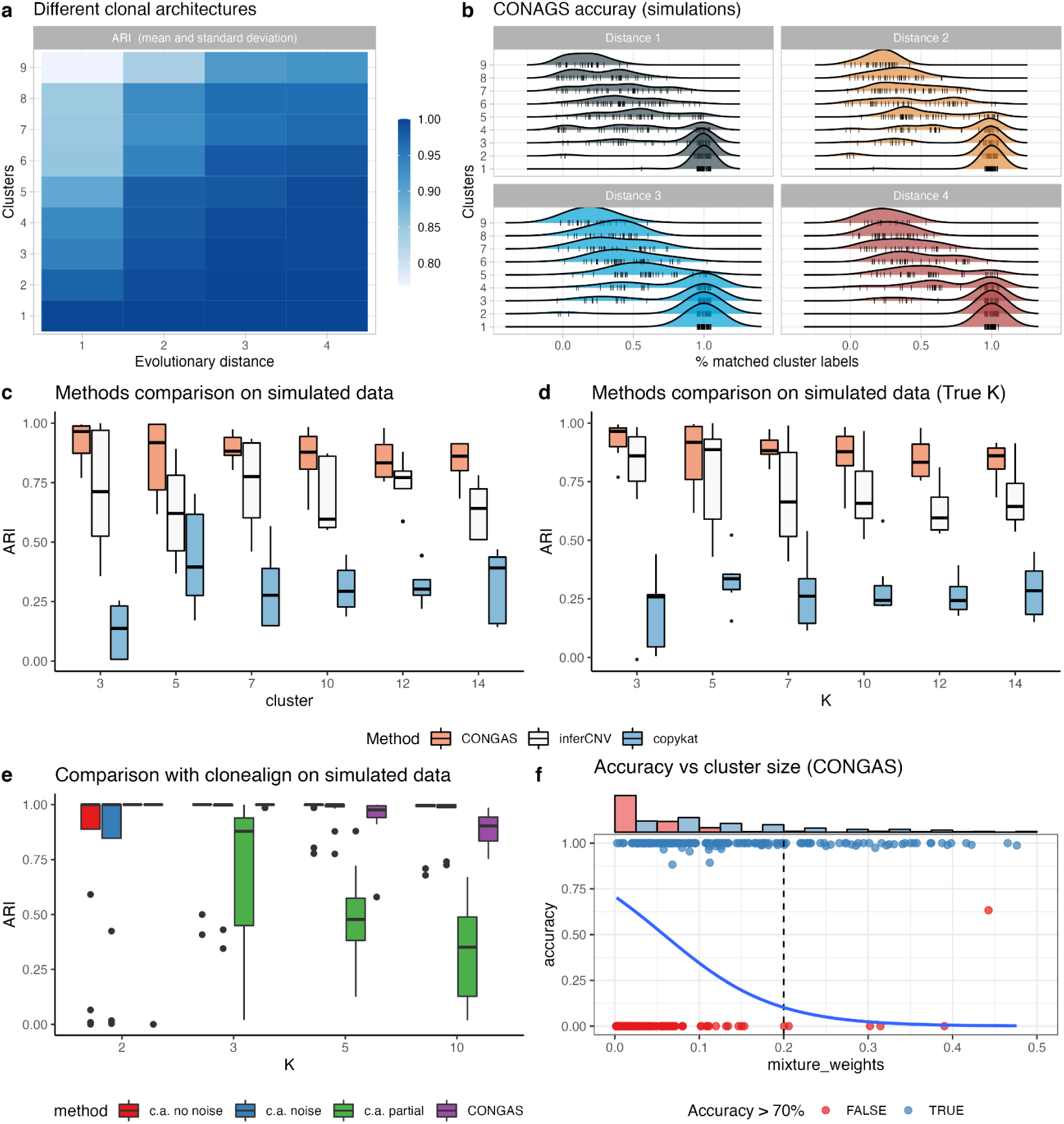
**a**. CONGAS synthetic tests with different subclonal architectures, obtained sampling clone trees with variable number of nodes. The degree of tumour heterogeneity is tuned by an evolutionary distance, which counts the number of CNAs that a subclone acquires, relative to its ancestor. The bulk input profile for CONGAS is generated by considering CNA segments from the most prevalent clone. We scan models with up to 9 clones, with distance ranging from 1 to 4. The performance is measured by using the adjusted rand index (ARI) between simulated and retrieved cell assignments. The heatmap reports the mean. **b**. Smoothed density for the percentage of cluster labels matched in every simulation, split by simulated tumour trees of increasing distance to mimic subclonal complexity **c**. CONGAS, inferCNV and copyKAT were run on a set of synthetic scRNA-dataset with 500 cells and a linear model for CNA effect on expression. Overall, CONGAS obtained the highest ARI score, and all other methods overestimated the true number of clusters in the data. **d**. The same simulations from panel (c) were re-clustered by cutting the dendrogram generated in output by copyKAT and inferCNV using the actual number of clusters. Despite the improvement in performance, especially for little *k*, CONGAS (unsupervised) continues to perform better than the other two tools. **e**. CONGAS and clonealign (supervised) were tested on a set of simulated dataset with the same generative process as in panel (b). For clonealign we tested three scenarios where we input the ground truth data (“no noise”), the ground truth data upon stochastically flipping subclonal CNAs (“noise”), a subset of the original subclones (“partial”). We observe that the ARI of both methods equates until many clones are present (>5); the performance of clonalign partial degrades largely. **f**. Performance of CONGAS in detecting clusters based on cellular proportion. Cases with accuracy above 70% are marked, showing the relation between cluster size and probability of detection. The blue line represents a logistic regression curve fit on the observed probability. The data is the same as panels (c,d).

CONGAS could also work with Negative Binomial overdispersed data, a violation of its Poisson model. Performance clearly increased for lower dispersion, plateauing for non-dispersed data (Supplementary Figure S4 and Supplementary Figure S5). We also tested how errors in the input segmentation affects deconvolution. Precisely, we generated subclonal CNAs that were shorter than the input bulk segments, so that only a percentage of genes mapping to a segment were showing a signal in RNA data (from 10% to 90% of mapped genes). This is another test-case where the assumptions of CONGAS are violated. We observed good performance when >40% of the genes that map to a segment are associated to the subclonal CNA (Supplementary Figure 4 and Supplementary Figure S6), which suggests that genotyping focal amplifications that involve a handful of genes might be hard, while larger CNAs are generally identifiable even with imperfect segmentation.

#### scRNA-based tools

We compared CONGAS against InferCNV and copyKAT, two popular CNA-calling methods for scRNA, using an independent single-cell RNA simulator to avoid biases (Zappia *et al*., 2017). We tested the performances with 500 cells from a variable number of subclones, assuming a linear model for the CNA-expression dependency. Overall, CONGAS obtained the highest ARI (always above 0.75 in all configurations), showing the ability to recover the real clusters in most cases. In general, CONGAS performance was particularly good in settings with <7 subclones, with clear difference to inferCNV. In those cases inferCNV showed a tendency to overestimate the real number of clusters by a factor of 2 (i.e., one false cluster for every true one), while CONGAS retrieves on average the exact number of subclones in the data. copyKAT showed slightly worse performance than inferCNV. From tests, we also observed that the probability of miscalling a cluster goes to zero as the size of the cluster increases, as corroborated by the fact that most of the clusters missed by CONGAS had less than 25 cells, and were therefore too small (<5% of the simulated cells) to detect (Figure 2c and 2f, Supplementary Materials). To avoid our conclusions being derived solely from using different model selection criteria, we compared the performance of inferCNV and copyKAT on the same dataset of simulations used previously, but this time giving the dendrogram cutting algorithm the true number of clusters. We indeed observed that the performances, especially for inferCNV, increase a lot for low *k*. Instead for *k* > 10 the ARI does not improve and in some cases (*k* = 12) decreases (Figure 2d, Supplementary Materials).

#### Clonealign

We compared CONGAS (unsupervised) against clonealign (supervised) using synthetic simulations and 3 possible inputs (Figure 2e, details in Supplementary Materials) in order to capture different qualities for the supervision set of clonalign. We considered *i)* the ideal input, when clonealign knows all the simulated clonal profiles (perfect clustering from scDNA-seq), *ii)* a noisy input, where we applied noise to the clonal profiles, simulating more realistic noisy scDNA-seq clustering and *iii)* a partial information, where only a subset of the real input profiles is given to clonealign. This last case simulates imperfect clustering from scDNA-seq (missing clones); this type of input could also mimic usage of a subclonal copy number caller from bulkDNA-seq (instead of scDNA-seq), where we call certainly fewer clusters than with a single-cell assay (Supplementary Materials).

We again generated assays with 500 cells using the same CNA model as the previous simulations. As expected, with prefect data clonealign has better ARI when the number of clones increases; in these cases since cluster size decreases with fixed number of cells, CONGAS is not able to separate well some clusters. Clonealign seems also very robust with respect to the adopted noise. On the other hand, when we simulate more realistic partial input profiles, the performance of clonealign decreases rapidly and proportionally to the number of clusters in the original data, and the performance of CONGAS is higher. Further comparison between the two tools are discussed below on real data collected from one triple-negative breast cancer.

### 3.2 Subclonal CNAs in a triple-negative breast xenograft

We used CONGAS to analyse a triple-negative breast cancer dataset generated with 10x technology; we use this case study to validate our method against single-cell low-pass DNA data, used initially for clonalign (Campbell *et al*., 2019) This dataset is the patient-derived xenograft SA501X2B collected from patient SA501, and has been used before to determine clone-specific phenotypic properties that associate with a complex clonal architecture, also validated by reproducing clonal dynamics over serial xenograft passages (Eirew *et al*., 2015). From low-pass whole-genome CNA calling, the authors estimated three genetically-distinct clones (prevalence 82.3%, 10.8%, and 6.9%); one clone sweeping in next engraftments.

To run CONGAS we retrieved the input genome segmentation from the largest clone identified in the original paper (82.3% of the cells). After retaining segments with at least 10 mapped genes and performing quality control, we retained *n* = 503 cells from which we could identify two of the three clones (Figure 3a). The signals identified by CONGAS are clear across multiple segments, with particular strength on chromosomes 15, 16 and 18 (two-sided Poisson test, *p* < 0. 001, Figure 3b). This is consistent with low-pass analysis (Campbell *et al*., 2019), validating our inference (Supplementary Figure S7, S8). Our analysis however does not detect the third subclone from the original analysis; this was explained by observing that subclonal CNAs defining that population contain <10 genes, and have been removed from data. We note however that this cluster is poorly supported also in (Campbell *et al*., 2019), which reports assignment uncertainty between the second largest clones and this population. Moreover, we tested if clonealign could have been used with a bulk whole-genome, instead of a low-pass single-cell one, to detect such cluster. In particular, we run the sublonal copy number caller ReMixT (McPherson *et al*., 2017) on bulk data from the primary tissue of SA501, and used its results as input for clonealign. Consistently with our analysis, in this case the tool was unable to discriminate the different populations (Supplementary Figure S7).

**Figure 3.**
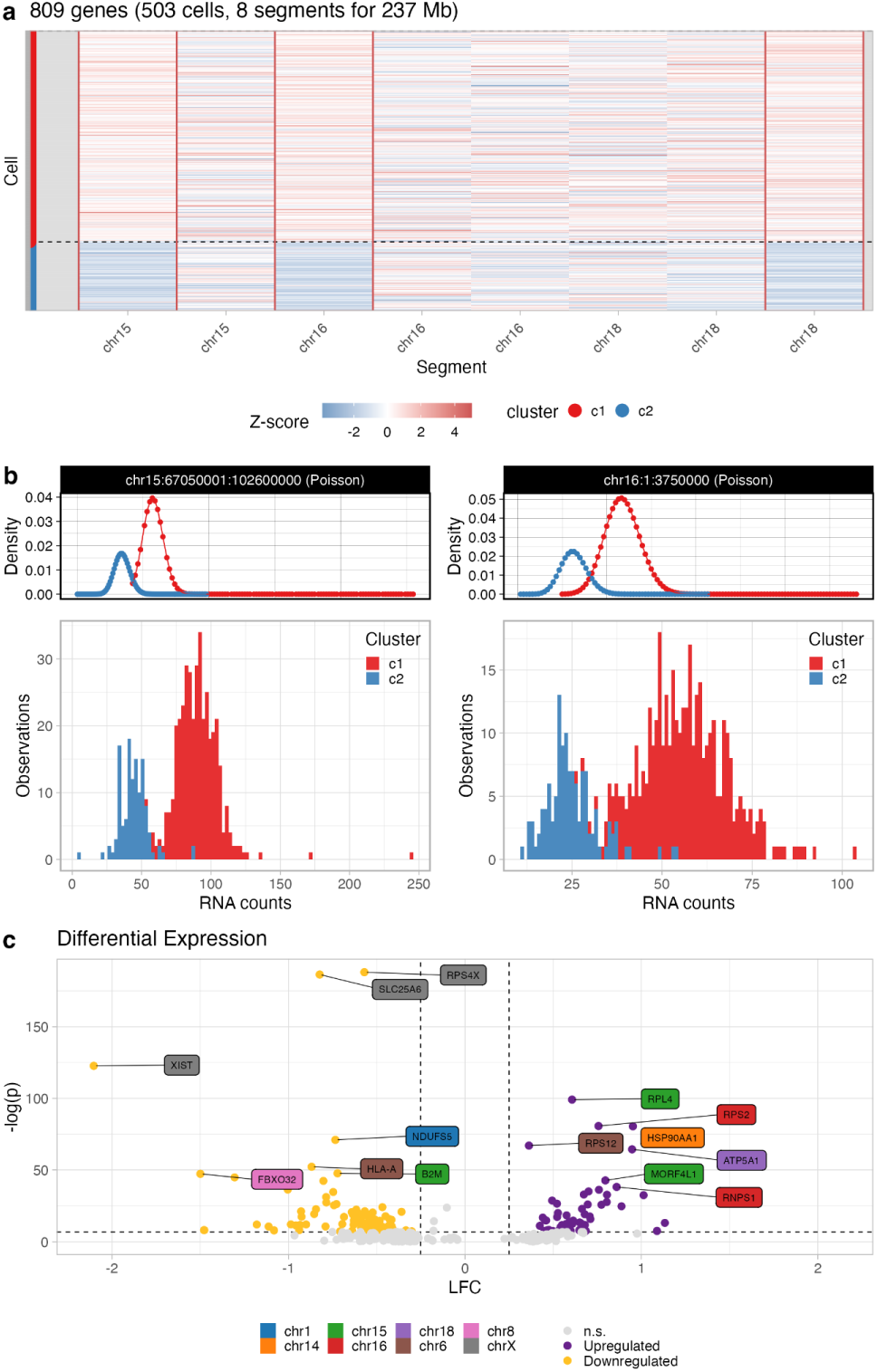
**a**. CONGAS analysis of *n* = 503 cells from a 10x assay from a triple-negative breast xenograft, where *k* = 2 populations are identified with 380 (~75%), and 123 cells (~25%). The heatmap shows input raw RNA counts (normalised per segment, with z-score) on chromosome 15, 16 and 18 where differences among CNAs are detected across the two subclones (red boxes). **b**. RNA transcripts count for the genes mapping to a segment on chromosome 15, and one on 16. The densities on top of the histograms are the Poisson mixtures inferred by CONGAS. **c**. Genome-wide clone-specific Differential Expression analysis highlights *n* = 212 dysregulated genes with adjusted *p* < 0. 01 and absolute log-fold change (LFC) >0.25 (up-regulation) or <0.25 (down-regulated); notice that some of those genes do not overlap with CNAs that characterise the populations.

The populations identified by CONGAS show significant differences in RNA counts (Figure 3b, Supplementary Figure S9): the largest subclone consists of *n* = 380 cells (~75%), and the smallest one of *n* = 123 (~25%). We also performed clone-specific differential expression analysis with the DeSeq2 (Love *et al*., 2014) and found (Figure 3c) 122 genes significantly upregulated or downregulated using a Wald test over negative binomial coefficients (adjusted *p* < 0. 001 via Benjamini-Hochberg correction), imposing absolute log fold change (LFC) >0.25 to determine the regulatory state (Figure 3c). Note that some of these genes do not overlap with CNAs, and therefore could only be marginally explained by genetic changes. Instead, they might be explained by more complex regulatory mechanisms indirectly linked to these, and other events. Library factors were also found quite variable across cells (Supplementary Figure S11).

We tested these data with inferCNV and copyKAT as well. Consistently with trends observed in simulations, while the true CNAs are identified even by these methods, the final number of clones is overestimated and spurious clusters are reported (Supplementary Materials, Supplementary Figure S12 and S13).

### 3.3 Tumour normal deconvolution in primary glioblastoma

We used CONGAS to analyse the glioblastoma Smart-Seq data released in (Patel *et al*., 2014). This dataset consists of *n* = 430 cells from five primary glioblastoma, from which we analysed patient MGH31 (*n* = 75 cells). MGH31 was chosen as it harbours subclones, according to both the original paper and a successive analysis (Fan *et al*., 2018). With this scRNA CONGAS was mainly challenged by i) the lack of an input CNA segmentation, and ii) the presence of normal cells in the assay (also known from the original analysis). To process this sample we have created a simple pipeline around CONGAS.

We have first developed a variational Hidden Markov Model to segment B-allele frequencies from germline single nucleotide polymorphisms called by scRNA (Supplementary Material). In this way we obtained segments with evident losses of heterozygosity, as well as large amplifications on chromosomes 7, 10, 13 and 14 (Supplementary Figure S14). In a first run (Figure 4b), CONGAS identifies *k* = 3 clusters from all cells (normal plus tumour); one of them (*n* = 10) lacked any CNA. The very same set of cells were classified as “normal” by a comparison with a healthy reference (Fan *et al*., 2018). We removed normal cells and re-run CONGAS on the remaining tumour cells, finding *k* = 3 distinct subclones (Figure 4a-c). These two-steps results were consistent with a solution with *k* = 4 clusters, obtained in the first run. Manual phylogenetic reconstruction after CONGAS suggested an early branching from an ancestor harbouring chromosome 7+ (amplification) and 10- (deletion). Clones then branched out: one sustained by 5+ (34% of cells), while a linear path described the evolution of two nested clones with increasing aneuploidy (with the largest subclone with 34% of cells), distinguished by 14- but harbouring the same deletion on chromosome 13 (Supplementary Figure S14).

**Figure 4.**
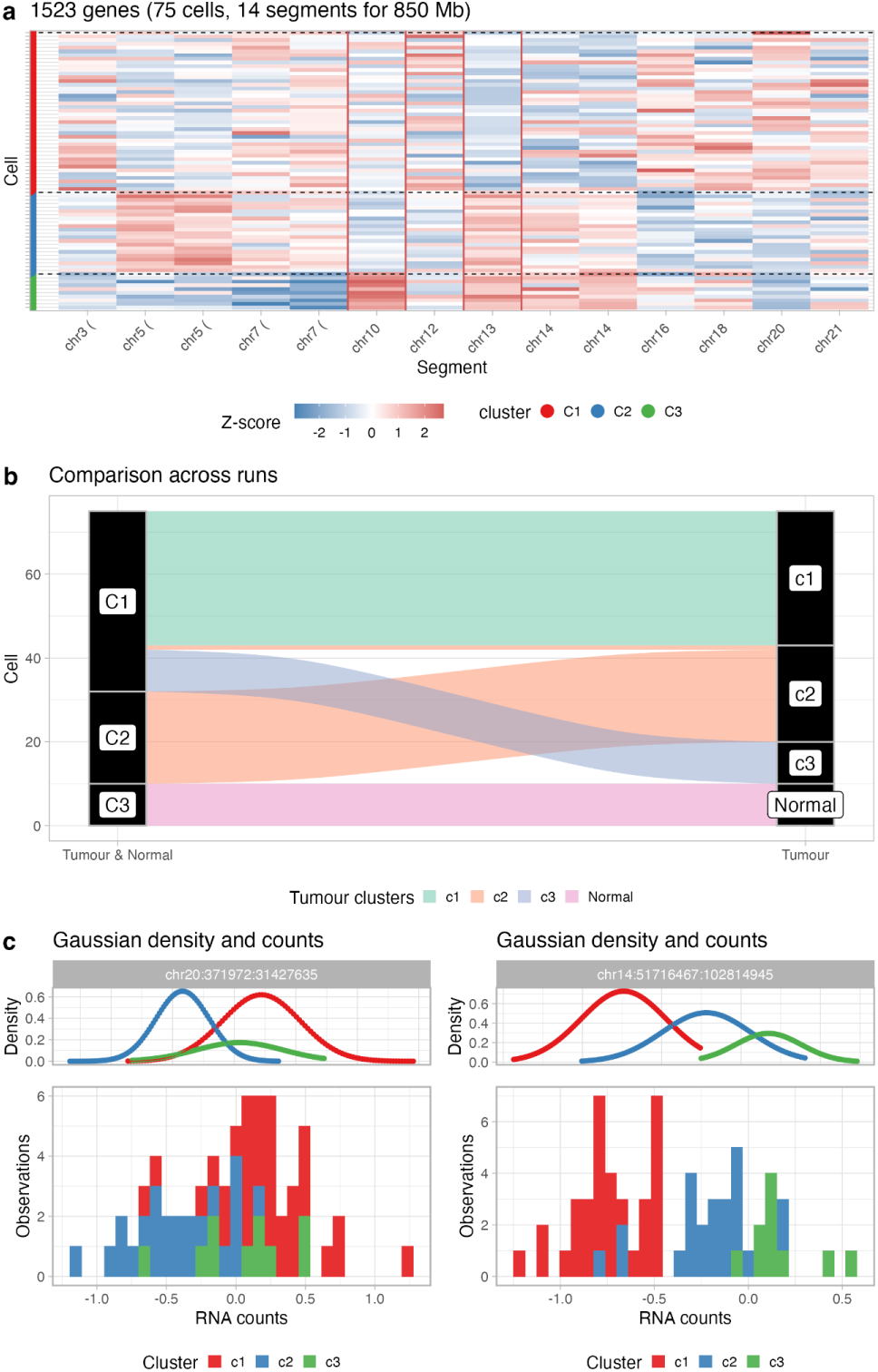
**a**. CONGAS two-steps analysis of *n* = 75 cells from a SmartSeq assay of a glioblastoma. The analysis first identifies normal cells in the sample, and then re-clusters tumour subclones; in the end *k* = 3 subclones are identified with 32, 23 and 10 cells. The heatmap shows input raw RNA counts (normalised per segment, with z-score) for a segmentation obtained directly from B-allele frequencies in RNA, and clusters from the first run (normal cells have no CNAs). **b**. Sankey plot of clustering assignments for the two runs. Cluster C3 from the first run are normal cells; tumour clusters are consistent across both steps of the analysis. **c**. RNA transcripts count for the genes mapping to a segment on chromosome 20, and one on 14. The densities on top of the histograms are the Gaussian mixtures inferred by CONGAS, here used instead of Poisson because data were normalised.

The DE analysis of these few cells was inconclusive due to the small number of sequenced cells (data not shown); nonetheless this 2-steps analysis shows how CONGAS can perform signal deconvolution in the presence of normal contamination of the input sample. This is interesting and consistent with the fact that the method can work without a reference normal expression.

We note that this data have been analysed also with inferCNV, honeyBADGER and CaSpER (Fan *et al*., 2018; Serin Harmanci *et al*., 2020; Patel *et al*., 2014). In all three cases, however, only two clones were found, one characterised by 5+, and another characterised by 13- and 14-, in substantial agreement with our analysis. However, our analysis is higher-resolution, since it splits the latter clone based on the presence or absence of 13- (Supplementary Figure S13).

### 3.3 Monosomy of chromosome 7 in hematopoietic cells

To show the versatility of CONGAS we have also analysed mixtures of non-cancer cells collected within one experiment associated with the Human Cell Atlas project (Rozenblatt-Rosen *et al*., 2017). In this case the dataset provides scRNA from hematopoietic stem and progenitor cells from the bone marrow of healthy donors and patients with bone marrow failure. We focused on one patient (patient 1) with severe aplastic anaemia that eventually transformed in myelodysplastic syndrome, and for which cytogenetic analyses revealed monosomy of chromosome 7, a condition that increases the risk of developing leukaemias (Zhao *et al*., 2017).

To analyse this data we pooled patient 1 together with one healthy donor, gathering *n* = 101 total cells (Supplementary Figure S15). This gives CONGAS both diploid cells (control, from the health patient), and cells with chromosome 7 deletion.

This dataset comes without a suitable input segmentation, so we used full chromosomes (arm-level segments) with a diploid prior. Aneuploid cells were clearly distinguished from diploid cells by CONGAS, which found *k* = 2 clusters. One, containing diploid cells from both patients, the other cells from patient 1 that are associated with monosomy of chromosome 7. Clone-specific differential expression performed as for the breast xenograft reported 99 genes differentially expressed at significance level *p* < 0. 01, and with absolute log-fold change >0.25. Interestingly, the top dysregulated genes were not expressed in the aneuploid chromosome, suggesting that an integrated study of transcriptomics and CNAs could lead to a better understanding of how these genomic events - which have considerable dimension - can alter cellular behaviour across different pathways and functional modules.

## 4 Discussion

In this paper we presented CONGAS, a Bayesian method to detect CNAs that can cluster single-cell RNA sequencing profiles, opening the way to study tumour subclonal composition at the single-cell copy number level. CONGAS requires inputs that can be generated by following a split design, leveraging both bulk and single-cell assays. In this way, the inference is easier and more precise compared to methods that call CNAs directly from scRNA. The method compares also against methods that assign single-cell RNA to subclonal CNA profiles, with the main advantage of being unsupervised. In this sense, input clonal CNAs are used to build a Bayesian prior to detect subclonal CNAS in single cells. In other approaches, instead, the clusters are pre-determined and cannot mutate during the cell-assignment process.

CONGAS also has other interesting features. First, it does not require a normal RNA reference from a matched tissue, or the presence of normal cells in the sample. This means that it can find subclones with different CNAs regardless of reference expression, a major advantage in organoids designs where we do not collect non-tumour cells (Vlachogiannis *et al*., 2018). Second, CONGAS reconciles copy number heterogeneity from RNA using a probabilistic model for cell assignment. Compared to callers that do not attempt clone detection or that separate calling from clustering, the advantage is that uncertainty is modelled in a unique framework, both for copy number estimation and clustering assignments. Third, the method uses a powerful probabilistic programming backend to scale to thousands of cells, overcoming computational limitations of other methods (Supplementary Figure S15 and S16).

CONGAS can be used to curate clonal evolution models (Caravagna *et al*., 2018, 2016), or to assess clone-specific phenotypic signatures at the RNA level. This mapping comes out as a byproduct of the integration of genetic copy number events together with RNA data. With CONGAS one detects CNA-associated subclones and their patterns of differential expression, a key step to study how selective pressures shape genotype and phenotype evolution in cancers (Caravagna *et al*., 2020). In addition, CONGAS is also able to correctly estimate the magnitude of subclonal copy-number events. Which together with the input segmentation obtained from bulk sequencing, allow the estimation of the subclonal karyotypic profiles. (Supplementary Figure S17, Supplementary Materials).

This work offers a complementary perspective to DNA-only methods, for which many single-cell CNA detection algorithms have been developed (Zaccaria and Raphael, 2020; Kuipers *et al*., 2020; Wang *et al*., 2018; Garvin *et al*., 2015; Macintyre *et al*., 2018). Working with DNA, these methods can infer a de novo segmentation of the tumour genome - i.e., without prior input segmentations - and in the future it will be key to integrate ideas at the core of these models together with RNA-genotyping methods such as CONGAS. Notably, in this work we also show - across multiple case studies - that we can determine clone-specific differentially expressed genes that can be explained only partially by copy numbers, pointing to complex non-trivial regulatory mechanisms that link genotype states with expression patterns. Our method provides a solid statistical framework to approach this type of investigation, which is crucial to determine disease clonal dynamics, as well as cell plasticity and patterns of drug response from the large wealth of single-cell data available nowadays.

## Authors contribution

SM, RB and GC conceptualised and created CONGAS, with support from LP and NC. SM and RB implemented the tool; SM ran synthetic tests, and collected data for the case studies with support from NC and RB. All authors analysed the data and interpreted the results. GC and SM drafted the manuscript, which all authors approved in final form.

