## Supplementary File 1 for "A Bayesian method to cluster single-cell RNA sequencing data using Copy Number Alterations"

### Supplementary Material

*Salvatore Milite, Riccardo Bergamin, Lucrezia Patruno, Nicola Calonaci, Giulio Caravagna*

### CONGAS

We discuss here our approach to genotype CNAs from single-cells RNA sequencing data. To be precise, while we generally refer to CONGAS as a single model, in reality the framework leverages a set of core ideas to create models with slightly different features, and that can analyse different types of data. The models are:

- (default) a Dirichlet finite mixture for RNA counts data;
- a Dirichlet finite mixture for log-normalised RNA data;
- a Dirichlet Process extension of both the above.

Also, the framework offers a simple Hidden Markov Model (HMM) to segment input single-cell data, which can be used to generate segmentations required by CONGAS if these are missing.

In next sections we discuss all models; plate notations are in Supplementary Figure S2.

**Modelling the relationship between CNAs and gene expression.** In order to recover distinct populations of cells that differ for the copy number of specific segments, we follow the idea of modelling the generative process of reads counting using a latent variable (Campbell et al. 2019). Instead of modelling the expression of a single gene, we use the aggregated read counts over a whole genome segment. This is why our model assumes a pre-existing segmentation of the genome. This segmentation is a fundamental part of the model, as it guarantees us a convenient way of treating the segments as independent statistical entities. As explained in the text, this extra assumption poses a minimal burden, considering the cost of generating the single-cell dataset.

The latent variable has a direct dependence on the copy number state, and it should explain the differential counts expression over a segment among clones. A simple but effective choice is to consider the segment expression to be linearly dependent on the total number of chromosomes; this is the same idea previously exploited in (Campbell et al. 2019). The other factors that contribute to segment expression and that have to be taken into account are the different depth at which each cell is sequenced, and the number of genes present in a segment.

Formally, we index with $i = (1, ..., I)$ the segments, with $n = (1, .., N)$ the number of cells, and we introduce a categorical latent variable for each cell assignment $\mathbf{Z} = [z_{nk}]$ such that

$$p(\mathbf{Z}|\boldsymbol{\pi}) = \prod_{n,k} \pi_k^{z_{nk}}$$

where $\mathbf{z_n}$ is a vector that evaluates to $1$ if cell $n$ belongs to cluster $k$ and $0$ otherwise, $\boldsymbol{\pi}$ is drawn from a Dirichlet distribution as in standard finite-mixture modelling (Caravagna et al. 2020).

We also indicate as $\theta_n$ a Gamma distributed latent variable which models the library size and name $\mu_i$ the number of genes in a segment $i$; this number is a constant that depends on the input segmentation. Our CNAs are modelled as continuous LogNormal distributions; we define $\mathbf{C} = [c_{ik}]$ such that $c_{ki} \sim \text{LogNormal}(m_{ki}, v_{ki})$ so to represent the copy number value for segment $i$, in cluster $k$.

In the default read-counts based mode in CONGAS, we describe the probability of the counts $\mathbf{Y} = [y_{ni}]$ of the cell $n$ in segment $i$ as

$$p(y_{ni}|\boldsymbol{\theta}, \boldsymbol{\mu}, \mathbf{C}, \mathbf{Z}) = \text{Pois}\left(\frac{\theta_n \cdot \mu_i \cdot \prod_{k=1}^{K} C_{ik}^{z_{nk}}}{\sum_{i=1}^{I} \prod_{k=1}^{K} C_{ik}^{z_{nk}}}\right)$$

and the full likelihood is obtained by assuming both cells and segments to be independent. This assumption is valid since we have an existing segmentation of the input genome. The model likelihood becomes

$$p(\mathbf{Y}|\boldsymbol{\theta}, \boldsymbol{\mu}, \mathbf{C}, \mathbf{Z}, \boldsymbol{\pi}) = \prod_{n=1}^{N} \prod_{i=1}^{I} p(y_{ni}|\boldsymbol{\theta}, \boldsymbol{\mu}, \mathbf{C}, \mathbf{Z}, \boldsymbol{\pi})$$

and a graphical representation of the model is in Supplementary Figure S2. Another way of thinking of the denominator in the formula is, given that all the effects are linear, as a matrix decomposition of the input. Note that here the denominator is omitted.

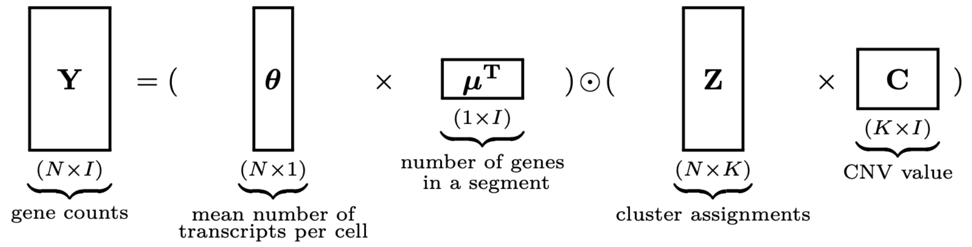

While this model does work with raw counts and accounts for both gene number and library size normalisation, the CONGAS framework also supports input single-cell data that are already normalised. This helps because often the only measurements availables are in units of transcripts per million reads, or alternative normalised measures. This is also useful if one wants to perform the inference using a custom normalisation method.

Therefore, in addition to the Poisson count-based model we have developed a model that works with Normal distributions. In this case we assume the input to be aggregate values (which are not anymore integers) of expression over all segments, which must be already normalised between cells and segments.

The segment expression $y_{ni}$ value in this alternative model formulation is now expressed as a Normal distribution; this model has likelihood

$$p(\mathbf{Y}|\mathbf{\Lambda}, \mathbf{C}, \mathbf{Z}) = \prod_{n=1}^{N} \prod_{i=1}^{I} \mathcal{N}(m_{ni} = \prod_{k} \mathbf{C}_{ik}^{z_{nk}}, \lambda_{ni} = \mathbf{\Lambda}_{\boldsymbol{i}})$$

Notably here we do not have any depence over library size and on the number of genes in a segment.

**Parameter estimation with variational inference.** Given our models we want to learn suitable values for the parameters, that for readability we indicate generally as $\mathbf{W}$ - so, in our case, $\mathbf{W} = (\mathbf{C}, \mathbf{Z}, \theta)$. For simplicity we drop any hyperparameter (our hyperpriors) or constants like the number of genes (μ) from the notation. Our goal, if we tackle the inference problem from a bayesian perspective, is to learn the posterior distribution of our parameters, namely

$$p(\mathbf{W} \mid \mathbf{Y}) = \frac{p(\mathbf{Y} \mid \mathbf{W})p(\mathbf{W})}{p(\mathbf{Y})}$$

Such distribution is generally analytically intractable (in particular the denominator, often called the evidence) and different methods for sampling or approximate inference have been developed. CONGAS is developed using the probabilistic programming language Pyro (Bingham et al. 2019) and exploits stochastic variational inference to get an approximation of the true posterior distribution (Blei, Kucukelbir, and McAuliffe 2017).

Briefly, the idea behind variational inference is to find a variational distribution $q(\mathbf{W})$ from a given distribution family $\mathcal{Q}$, so that $q(\cdot)$ can approximate the real posterior $p(\mathbf{W} \mid \mathbf{Y})$. Once the distributions family is fixed the problem of learning a posterior can be reframed as an optimisation task. More in detail we want to solve the optimization problem

$$q^*(\mathbf{W}) = \underset{q(\mathbf{W})\in\mathcal{Q}}{\arg\min}\ \mathrm{KL}(q(\mathbf{W})\|p(\mathbf{W}\mid\mathbf{Y}))$$

where $KL(\cdot)$ is the Kullback-Leibler (KL) divergence among probability distributions (Blei, Kucukelbir, and McAuliffe 2017).

This problem however involves calculating the posterior (and thus the evidence), in the above formulation, and needs to be rephrased with a more tractable objective function. Rewriting the equation above using simple laws of probability leads to

$$\mathrm{KL}(q(\mathbf{W})\|p(\mathbf{W}\mid\mathbf{Y})) = \mathbb{E}[\log q(\mathbf{W})] - \mathbb{E}[\log p(\mathbf{W},\mathbf{Y})] + \log p(\mathbf{Y})$$

By noting that the marginal log-likelihood $\log(\mathbf{Y})$ is a constant term given the data, we can perform optimization directly on the remaining terms.

$$\mathrm{ELBO}(q) = \mathbb{E}[\log p(\mathbf{W},\mathbf{Y})] - \mathbb{E}[\log q(\mathbf{W})]$$

This quantity is known as the Evidence Lower Bound (ELBO), as it becomes exactly equal to the evidence when our variational posterior is exactly equal to the real posterior, otherwise this quantity is always a lower bound for $\log(\mathbf{Y})$, since the KL between two distributions is always positive by definition.

In particular, our inference is done performing gradient descent optimization over the ELBO, that here we have rewrote in a form that highlights the contribution of the likelihood and the KL between the original prior and the variational posterior.

$$\mathrm{ELBO}(q) = \mathbb{E}[\log p(\mathbf{Y}\mid\mathbf{W})] - \mathrm{KL}(q(\mathbf{W})\|p(\mathbf{W}))$$

As we have discussed above, this is equivalent to minimising the KL divergence between the variational and the actual posterior distribution. In this framework our latent variables $q_\phi(\mathbf{W})$ are parameterized by a set of variational parameters $\phi$ and our goal is to learn those parameters by maximising the negative ELBO. Taking the gradient from the previous equation and expliciting the dependence on the variational parameters $\phi$

$$\nabla_\phi \mathrm{ELBO} = \nabla_\phi \mathbb{E}_{q_\phi(\mathbf{W})}\left[\log p(\mathbf{Y},\mathbf{W}) - \log q_\phi(\mathbf{W})\right]$$

We used the reparametrization trick to obtain a low variance Monte Carlo estimate of this gradient, and the tool can choose to calculate the gradient estimate over a minibatch of observations. The minibatch still provides an unbiased estimation of the gradient and can give a huge speedup over big datasets. We used Adam (Kingma and Ba 2017) as the optimizer of our choice throughout the whole paper, nevertheless the user can use any optimizer present in PyTorch (Paszke et al. 2019) or define a custom one.

The tool also gives the possibility to perform Maximum A Posteriori (MAP) inference, using the same mechanism.

$$\hat{\mathbf{W}}_{\mathrm{MAP}}(\mathbf{Y}) = \underset{\mathbf{W}}{\operatorname{argmax}}\, p(\mathbf{Y} \mid \mathbf{W}) p(\mathbf{W})$$

Note that the latent variable describing the library size dependence $\boldsymbol{\theta}$ is always learned using MAP inference, while for the other variables the user can choose between a full Bayesian inference, or MAP.

**Model and prior distributions.** The Bayesian setting gives us the opportunity to integrate some pre-existing information directly into the model. A way to guide solutions toward a meaningful direction is to assume the prior distribution of CNV values (the $c_{ij}$ in our model; see also Figure 2b) to be centred around the copy-number values obtained from bulkDNA-seq analysis.

More in detail, the model joint distribution can be factorised as:

$$p(\mathbf{Y}, \mathbf{Z}, \mathbf{C}, \theta, \boldsymbol{\pi}) = p(\mathbf{Y}|\mathbf{Z}, \mathbf{C}, \theta, \boldsymbol{\pi}) p(\mathbf{Z}, \mathbf{C}, \theta, \boldsymbol{\pi})$$

$$= p(\mathbf{Y}|\mathbf{Z}, \mathbf{C}, \theta, \boldsymbol{\pi}) p(\mathbf{Z}|\boldsymbol{\pi}) p(\boldsymbol{\pi}) \prod_{ik} p(c_{ik}) \prod_{n} p(\theta_n)$$

In the variational framework our latent variables are approximated as variational distributions distributions $q(\mathbf{Z}, \mathbf{C}, \theta, \boldsymbol{\pi})$, supposed to be independent and factorizable. The prior distributions for our latent variables are:

- $p(c_{ik}) \sim \mathrm{LogNorm}(m_{ik}, v)$, where $m_{ik}$ is the CNV value from bulkDNA-seq and the variance $v$ is chosen by the user to govern how far we think the actual CNV values are from the ones inferred by bulkDNA-seq, we use a default of 0.5.
- $p(\theta_n) \sim \mathrm{Gamma}(e_s, e_r)$, here $\theta$ distribution can be roughly estimated from the data; however, even large scarcely informative priors tend to work well in most of the cases. Default values are $e_s = 3$ and $e_r = 1$
- $p(\boldsymbol{\pi}) \sim \mathrm{Dirichlet}(\mathbf{r})$, the user can input is prior over the cluster distributions, by default all cluster are a priori assumed to have equal proportions (i.e. $r_k = 1/K$)

Variational distributions are chosen to be from the same family as the priors.

In the Gaussian mixture model, $q(c_{ik})$ prior is still distributed as a LogNormal, with the same parameter as before, while for the variance prior $\lambda_i \sim \mathrm{Uniform}(a, b)$.

For the first step of optimization $\mathbf{C}$ is initialised using k-means clustering, the library size factor is initialised as

$$\boldsymbol{\theta_n} = \frac{\mathbf{x_n}}{(\boldsymbol{\chi}^T \boldsymbol{\mu})}$$

where $\boldsymbol{\chi}$ is the CNV value obtained from the input bulk segmentation.

The decision to use a continuous latent parameter $q(c_{ik}) \sim LogNorm$ for the copy-number values instead of a more common discrete variable is due to two main considerations:

- the perfect linearity of copy-number values and expression is an imperfect assumption, real data is often complex and noisy, and a continuous latent variable is more flexible;

- our inference method is based on gradient propagation through stochastic functions, one of the hardest distribution to deal with in this framework is indeed the categorical one, which often inflate the variance of the gradient estimator (Jang, Gu, and Poole 2016; Schulman et al. 2015).

**Model-selection.** All these formulations assume the number of clusters $K$ to be a constant. To pick the optimal number of clones we first select a set of candidate $K$s and fit a separate model for each one, then we perform model selection. In the our packages the information criteria currently implement are:

- the Bayesian Information Criterion (BIC) $\mathrm{BIC} = k \ln(n) - 2 \ln(\widehat{L})$, where $\hat{L}$ is the likelihood of the model;
- the Akaike Information Criterion (AIC) $\mathrm{AIC} = 2k - 2\ln(\hat{L})$;
- the Integrated Classification Likelihood (ICL), based on the BIC, which is

  $$\mathrm{ICL} = \mathrm{BIC} + \mathrm{H}(\mathbf{Z})$$

  where $\mathbf{Z}$ are latent variables and $\mathcal{H}$ their entropy

  $$\mathcal{H}(X) = -\sum_{i=1}^{n} \mathrm{p}\left(x_i\right) \log \mathrm{p}\left(x_i\right) \ .$$

Where not explicitly indicated for our in silico and in vivo analysis the criterion of choice was always the BIC.

Total number of cells and segments has to be taken into account, as for some Smart-Seq runs with a small number of cells the BIC and ICL regularisation might be too strong. In those cases filtering segments or using other selection criteria can help to obtain meaningful results. On the other hand when the number of cells is higher than 200/300 one can normalise the likelihood by the number of segments to avoid overclustering.

**Non-parametric extension.** Given the importance of selecting the correct number of clusters in a finite mixture model, we also adopted a non-parametric formulation of CONGAS. This is expressed as a stick-breaking formulation of the Dirichlet Process (Ferguson 1973).

The stick breaking model in our case can be seen as a semi-parametric way of choosing the optimal number of clusters along with their weights. Note that the Dirichlet Process is defined for an infinite number of clusters, what is commonly done in practice is to set a high number of clusters to approximate this behaviour.

The generative process in our setting has these steps (Ishwaran and James 2001):

- Draw $\beta_k \sim \mathrm{Beta}(1, \alpha)$
- Draw $c_{ik} \sim \mathrm{LogNormal}(m_{ki}, v_{ki})$
- The mixture weights $\pi$ are obtained as

$$\pi_k(\beta_{1:T} = \beta_k \prod_{j<i}(1 - \beta_j))$$

- For each $n \in 1, ..., N$ and $i \in 1, ..., I$, draw $z_n \sim \pi_k$ and $x_{ni}(c_{ik})$

Where we omitted all the variables that do not depend explicitly on the clusters $K$. The $\alpha$ is a hyperparameter that controls our prior beliefs on the number of clusters present in the data, i.e. higher $\alpha$ will penalise a solution with more clusters and vice versa. Learning good values for $\alpha$ is fundamental in a noisy setting like scRNA-seq, and different metrics have been described for such optimization problems (Blei and Jordan 2006). Nevertheless, throughout the paper we preferred to run different instances of the finite mixture model and then do model selection as explained above. This is due both to the difficulties in accurately optimising $\alpha$ and in the increased understanding of the data one can get from having all the models fitted.

### Performance of CONGAS

A batch of datasets for performance assessment was generated using CONGAS's generative model. First, the human chromosomes (reference hg38) were divided into segments. For the segmentation we first set an approximate minimum distance for the segments (100 Megabases) and then divide the total length of each chromosome. This process gives us the maximum number of possible segments for a chromosome, let us call it $m_{chr}$. We sampled the effective number of segments $i_{chr}$ as an integer ranging from 1 to $m_{chr}$, with equal probability. Breakpoints were then calculated by dividing each chromosome by $i_{chr}$ and then randomising the endpoints of each segment adding an error value $e_i \sim \mathrm{Unif}(a, -a)$, where $a$ is equal to 10% of the segment length. The total number of genes for each segment were sampled from a negative binomial distribution with size $6$ and mean equal to segment length divided by a constant, in our case $1e6$.

Independently, a tree with a number of clones $K$ and a distance among clones of $D$ is constructed. The distance here is just the Hamming distance between segments, i.e. the number of different segments between two clones. The tree, given the number of clones, is set up by iteratively adding each new node to it and choosing its parent randomly from the tree leaves (with equal probability). To simplify both simulation and interpretation of the results each branch of the tree has the same length so that each child diverges from its parent for the same number of CN events. Using this information we sampled counts for CONGAS.

For the first test we simulated trees with a number of clones ranging from 1 to 9 and distance from 1 to 4, with 50 replicates for each combination. The clone proportions were randomly sampled from a Dirichlet with concentration parameters of $\alpha_k = 1/K$.

For the second test we changed the generative model to obtain overdispersion in counts data sampled from a negative binomial with parametrization

$$\Pr(X = k) = \binom{y + \zeta - 1}{y} \left(\frac{m}{m + \zeta}\right)^{y} \left(\frac{\zeta}{m + \zeta}\right).$$

Here the mean is equal to $m$, $\zeta$ is the size and the variance is

$$\mathrm{Var}[X] = m + \frac{m^2}{\zeta}$$

so that our likelihood becomes

$$p\left(y_i^n \mid \theta, \boldsymbol{\mu}, \mathbf{C}, \boldsymbol{\beta}, \mathbf{z}\right) \sim \mathrm{NegBin}\left(m = \theta_n \cdot \mu_i \cdot C\left[z^n, i\right] \cdot \exp^{\mathbf{w}_n^{\mathrm{T}} \beta_i}, size = \zeta\right)$$

We tested size values $\{2, 4, 8, 16, 32, 50, 100, 150, 200\}$, runned 50 repetitions for each and fixed the number of clones to $2$, sampled the abundance of each clone in the range $[0.7, 0.3$, and set the distance to $3$.

In the third test we analysed cases in which the input bulk segmentation does not faithfully represent the real subclonal segmentation. In the standard CONGAS model, for every gene that maps to a segment, the linear relation between segment ploidy and RNA counts always holds. In this test we simulated a violation of this assumption, allowing that in the same segment just a percentage $\gamma$ of the total number of genes is affected by the copy number state. For this test we evaluated five setups, where $\gamma$ starts from 20%, reaches 100% (like in the first test), with steps of 20%. We again performed 50 repetitions for each value of the parameters. To focus on the role of this assumption, we adopted simple models with just two segments, one of which harbouring the same copy-number among clones. Here the number of clones, abundance and distance were fixed to $2$, to the range $[0.7, 0.3]$ and to $1$.

We assessed CONGAS performance by comparing the inferred with the ground truth clusters. In the case of simulated data, for any number of clones and repetitions we matched a cluster inferred by CONGAS with a ground truth cluster on the basis of the overlaps of cell assignments. For any couple of clusters we computed the euclidean distance between the copy numbers of all the segments. We did the same test also for the triple-negative breast xenograft dataset. We reported the histogram distribution of all the euclidean distances for a simulated dataset with two clusters and for the triple-negative breast xenograft dataset in Supplementary Figure S17a-c. For the cases with two clusters we also assessed the ability of the tool to distinguish two subpopulations on the basis of amplifications and losses. For all the segments we computed the difference between the copy number of the two clusters. We reported the trend of the copy number difference for the inferred and ground truth clusters in Supplementary Figure S17b,d for the same simulated example with two clusters, and for the triple-negative breast xenograft dataset.

For simulated datasets results show that CONGAS is able to infer the relationship between the two clusters (Supplementary Figure S17b), correctly identifying segments where both clusters have the same copy number (i.e., the difference is 0) and segments characterised by different copy number states. In detail, in the reported example there are three segments where the first cluster has a lower CN value with respect to the second one, and we show that CONGAS identifies this trend for all of them. This is reflected also on the Euclidean distance (Supplementary Figure S17a), where the difference between ground truth and inferred copy number state of each segment is ≈ 0.

For the triple-negative breast xenograft dataset, Supplementary Figure S17c,d show the corresponding results. For three segments the difference between ground truth and inferred trends is higher than 0.5. However, we can see that these segments are characterised by a low number of genes mapped onto them, which indicates that the low amount of signal affected the inference performed by CONGAS.Overall we can see that the trend between the two clusters is recovered, as most segments display a difference lower than 0.5 (Supplementary Figure 17d). This is reflected also on the Euclidean distance (Supplementary Figure 17c), where, with the exception of specific outliers, most of the values are ≈ 0.

#### In-silico comparison against competitor tools

We compared CONGAS to other popular CNA-inference methods for scRNA data, restricting to those with inputs comparable to our approach. In particular, we tested inferCNV (Patel et al. 2014) and copyKAT (Gao et al. 2021) as the other tools were using additional data (CaSpER, for instance, needs allelic frequency values) or were not well documented and reproducible.

To perform an unbiased comparison, we avoided using the generative model of CONGAS to create input datasets, and switched to the popular Splatter tool for scRNA simulation (Zappia, Phipson, and Oshlack 2017). We generated synthetic datasets consisting of 500 cells; Splatter parameters were estimated using the PDX breast-cancer dataset as a reference (with the "splatEstimate" function). Counts were obtained by using the "splatSimulate" routine with default parameters, we simulated 15000 genes for each run. We then randomly assigned the synthetic genes generated by Splatter to known human genes in chromosomes 1-22. For each run a random clonal tree was constructed as described in the previous section, and used to draw the mixing proportion for each node from a Dirichlet distribution with uniform concentrations (1/K, where K is the number of clusters), and assigned the cells to each clone with probability proportional to those mixing factors.

We assumed a perfect linear relationship between CN and expression, if we call $c_{ik}$ the CNA value of a given subclone $k$ in segment $i$, and $e_i$ the expression value for that segment in a normal diploid cell, than the final value of expression for that segment in the tumour subclone is $e_i c_{ik}/2$ (e.g., if the cell is tetraploid in a segment, then all genes in that locus have double the expression relative to baseline).

To let inferCNV and copyKAT operate in a fair comparison, one of the clusters was always composed of diploid cells (baseline Splatter signal). While CONGAS and inferCNV have an automatic way to determine the final number of clusters (CONGAS uses a mixture model, while inferCNV uses random trees after inferring CNAs), copyKAT returns only a tree built on the distances between cells and a label for diploid/aneuploid. As our comparison in this specific case is focused on inferring the correct number of clusters, we had to use the dynamicTreeCut R package to cut the histogram and obtain copyKAT clusters (Langfelder, Zhang, and Horvath 2008). dynamicTreeCut provides two different cutting algorithms, one using only the dendrogram, one integrates also the distance matrix. We used the latter approach as it works generally better and uses all the information available.

We tested CONGAS, inferCNV and copyKAT on a number of subclones ranging from 3 to 14. We performed 10 repetitions for each number of clusters and calculated the ARI to evaluate the performances of the different methods. CONGAS performs slightly better than inferCNV, which in turn is better than copyKAT (Figure 2, Main Text). We note that copyKAT does not have an automatic method for inferring the clusters, and the choice of our cutting strategy is arbitrary, and may be suboptimal. There is however an evident difference in the trends of CONGAS and inferCNV. CONGAS performs generally better with a low number of clusters, and the performance slightly decreases if the number of clusters increases. On the other hand, inferCNV shows the exact opposite behaviour, calling an average number of 4.7 and 6.5 clusters when the original K is respectively 3 and 5 (overestimating $K$, while CONGAS calls on average 2.8 and 5 clusters) and improving while K goes to 14. As K goes over 10 however, both tools identify an average of almost 8.5 clusters while still achieving good performances, this can be due to the high number of clusters with extremely low abundance that results from the Dirichlet sampling and the fact that we have simulated 500 cells. Overall, tests show how CONGAS is an optimal tool when the focus is retrieving the optimal number of clones, in scenarios similar to real data (i.e., with fewer than 7 clusters in the data). From the simulations we obtained the accuracy of the inferred CONGAS clusters against the simulated mixing proportions. As expected CONGAS accuracy is strongly correlated with cluster size, in particular the rate of missed clusters decreases sharply for larger clusters. Most miscalled clusters are in the 0-5% range, while for greater values CONGAS rate of true positives exceeds false negatives. Overall, the rate of missed clusters is extremely low if the cluster is larger than 15% of the sequenced cells.

These results provide guidance on the expected probability of correctly identifying a clone, based on its abundance. It is important to note, however, that those conclusions are based on simulations with 500 cells, and one expects the resolution of the method to scale with the sample size (based on BIC and ICL formulas). Importantly as we have tested a big number of cells with a relatively low number of segments we runned CONGAS with the option *normalize_by_segs=TRUE* as it helps the regularisation when the number of cells is high or the signal is particularly variable.

We also evaluated the performance of CONGAS against inferCNV and copyKAT on simulated data, with a fixed number of clusters corresponding to the true number of subclones. The simulated data is the same used for the unsupervised test but this time we cut the resulting trees using the correct number of clusters with the R function *cutree* from the *dendextend (Galili 2015)* package and reassigned cells accordingly. The results show that CONGAS outperforms both copyKAT and inferCNV, for all the tested numbers of subclones (Figure 2d). Nevertheless, the performances, particularly of inferCNV, increase significantly, especially for low $k$. Instead for large $k$ the ARI does not improve and in some cases decreases. This is mainly due to the higher frequency of small clusters that are very difficult to call for the tools and therefore not clearly visible in the tree. These clusters however, even if they do not generate well separable splits in the dendrogram, are still counted in the theoretical $k$ and therefore force the tool to further split monophyletic clusters. This result is even more relevant given that model selection is a fundamental part of the CONGAS framework. Thus comparing the performance of CONGAS against copyKAT and inferCNV with a fixed number of clusters provides biassed comparisons that do not take into account the model selection included in our model.

### In silico comparison against clonealign

We evaluated CONGAS against clonealign on a set of simulated examples. For consistency with other tests, we used simulated data generated in the same way as in section *In-silico comparison against competitor tools* section. Specifically, we tested a variable number of clusters (K = {2,3,5,10}) in order to cover a plausible range of clones. For each value of K, 8 datasets of 500 cells each were generated. As clonealign is designed to spot subclones in the tumour population we did not introduce any healthy diploid population in this batch of simulations, i.e. the purity was always 1.

We generate 3 use cases, based on the input we used for clonealign:

1. the true copy number profiles for all the subclones present in the sample;
2. like in point 1, but where we randomly shuffled the copy number value for each segment by a factor (-1,+1) with probability 20%. This was to simulate an incorrect call stemming from the scDNA-seq copy number caller;
3. a subset of the true copy number profiles from point 1. We simulate a dropout mechanism: we first drawn $\mathbf{x} \sim Binom(0.2, K-1)$ with $\mathbf{x} = x_1, , x_{K-1}$ where K is the number of subclones for a given sample and then we eliminated any subclone from the clonealign input matrix if $x_{(k+1)} = 1$.

Note that the sampling in case (3) is restricted to K-1 subclones, therefore we always keep the first cluster to avoid pathological settings. Comments on this test are in the Main Text.

#### Triple-negative breast xenograft analysis

Data was obtained from (Campbell et al. 2019), in the form of a count matrix for which we implemented some basic preprocessing. We removed from the count matrix all those genes expressed in less than 5% of the cells, furthermore we filtered the 5% most expressed genes, as their fluctuation could influence too heavily the total segment counts. All the cells with less than 3000 expressed genes were also removed. We calculated the total number of counts in the segment as the sum of the gene counts completely overlapping with a genomic segment.

The input segmentation for CONGAS was again retrieved from (Campbell et al. 2019), even if in the original papers the authors used low-pass single-cell DNA sequencing (instead of bulk, as we assume in CONGAS), and clustered cells defined by similar copy-number events. To obtain a unique input profile for CONGAS we considered the CNC profile of clone A from the original analysis. This is a good surrogate of putative values that we could obtain from a DNA sequencing assay, given that the abundance of this clone is above 80%.

To analyse these data we used a two-steps procedure We first runned the inference with default CONGAS parameters, using a learning rate of 0.01 for 800 inference steps. Posterior probabilities were computed with another 100 steps, using a learning rate of 0.05. In this first run the latent distribution over the CNA values was learned by MAP inference. In this way we set normalisation factors and CNV latent variables near to good solutions, as learning the full model together is usually less stable and shows marked multimodality.

In the second step we rerun the same model forcing a full Bayesian setting, where we now can learn the full distribution of our latent variables. In this case we focused on learning mean and variances of the LogNormal densities that model CNA values, while fixing the other parameters identified in the previous run. We obtained the final clustering assignments after filtering the clusters with an abundance of less than 3%.

Differential expression was performed using Seurat (Stuart et al. 2019) and DESeq2 (Love, Huber, and Anders 2014). Concordance between our clustering assignment and the labels of clonealign was quantified using Adjusted Rand Index (ARI), the corrected-for-chance version of the Rand index.

We run inferCNV and CONGAS on the same dataset, with the same preprocessing filters used for CONGAS. For InferNCV we used default parameters except for *cutoff* which we set at 0.1 (as suggested by the authors when dealing with 10x data) and *analysis_mode* to "subclusters", to also identify subclonal CNVs. We used the standard settings by acquiring a matched healthy dataset from GTEx (Supplementary Figure S12a-b), selecting tissue from breast (GTEx Consortium 2020). As inferCNV is theoretically able to run also without reference normal - it still detects the subclonal structure without explicit copy number values - we also run it without reference (Supplementary Figure S12c-d).

The performance of the reference-normalised algorithm and the reference-free are relatively similar, at least for what it concerns subclonal deconvolution. Clearly, without whole-genome sequencing or healthy RNA references it is impossible to identify the magnitude of the copy-number event from transcript counts. With these data where we know the ground truth (2 clones in total), inferCNV splits the correct subclone into two. Furthermore, the rest of the tumour is in turn divided into 5 subclones. InferCNV therefore confirms the impression it had with the simulations, i.e., that it tends to overestimate the number of subclones in the data.

We run copyKAT without diploid cells. Inference was done with default parameters. In this case, as in the simulations, to cut the tree we used the hybrid algorithm of DynamicTreeCut (Supplementary Figure S13c-d). Also in this dataset the determination of the correct number of clusters remains challenging, while the main CNAs are identified by all the methods. The combination of DynamicTreeCut and copyKAT however overfits the data with 4 clones identified, but to a lesser extent than inferCNV.

#### Glioblastoma data analysis

Input Smart-Seq scRNA data were obtained from (Fan et al. 2018). The original analysis does not provide an input segmentation for CONGAS. We developed a simple Hidden Markov Model (HMM) to segment exomes directly from RNA counts, and generate the missing segmentation.

**HMM definition.** Without assuming normalised healthy data, we can segment the genome by using allele frequencies. This approach works better with the Smart-Seq protocol than with the 10x one as it covers the whole gene instead of just the 3'/5' end. We identified heterozygous single nucleotide polymorphisms (SNPs) and calculated the ratio between counts for the major and minor alleles, deriving the minor allele frequency (MAF). In a healthy genome this is 0.5, if we disregard Binomial observational noise.

Our input data consists of mostly cancer cells, and MAFs are therefore no longer necessarily distributed around 0.5. Of course, the resolution of MAFs is limited with single-cell data, and we cannot expect to distinguish all segments breakpoints, or segment perfectly all the genome

The HMM is developed in Pyro and is available in the CONGAS package. The HMM has 6 hidden states; we consider CN values above 5 unlikely, at least for the purpose of segmentation. Furthermore, our ability to identify different states from MAFs deteriorates with very high CN. Priors on hidden states are described by the following matrix

$$\begin{pmatrix} 1-5t & t & t & t & t & t \\ t & 1-5t & t & t & t & t \\ t & t & 1-5t & t & t & t \\ t & t & t & 1-5t & t & t \\ t & t & t & t & 1-5t & t \\ t & t & t & t & t & 1-5t \end{pmatrix}$$

Here $t > 0$ is the propensity to change the internal HMM state, and is the same across states. In order to perform some filtering on the data, we set a small value for $t$, with default $10^{-6}$. The emission probabilities for MAFs are instead Beta distributions ranging in $[0, 1]$and are parametrized to be distributed around theoretical MAF values for each ploidy state. Note that the first state identifies LOHs, which are not automatically associated with the real number of copies.

To infer the HMM parameters we use SVI, with MAP inference for the transition matrix and the initial state vector. We calculate the posteriors for the state assignments. The emission probabilities, on the other hand, remain fixed and are not learned. We show runs of this HMM with the glioblastoma Smart-Seq in Supplementary Figure S14.

**Healthy cells contamination and subclonal detection.** The data for obtaining the MAF and the TPM normalised count matrices were taken from (Fan et al. 2018) .

To avoid RNA-editing the snps have been restricted to those present in the ExAC database with a frequency greater than 10% (Lek et al. 2016). Our segmenter works with the MAF for the whole tissue, so the first thing we did was to add the reference and alternative allele values for all cells and create a pseudo-bulk. Sites with coverage less than 20 reads were discarded. Our HMM segmenter was run on those variants with a t of 1e-8 and a median filtering window of 25 sites. After a manual examination we decided to collapse all the states different from a putative LOH (i.e. those different from 1), as the variance of the MAF was too high to confidently call the other CN states.

We then runned CONGAS over this segmentation, and obtained an ideal number of clusters equal to 3. Among those we could clearly identify one (cluster number 3) consisting of normal cells, as it was lacking any LOHs. We then excluded the cells belonging to that cluster, recalculated the pseudo-bulk MAF, filtered sites with less than 30 reads and rerun the HMM segmenter. Also in this case states with a theoretical MAF value too close to each other were collapsed together, in particular 2 with 6 and 4 with 5. This new putative segmentation was given as an input for a new CONGAS run. The tool was able to identify 3 putative clones characterised by specific chromosome alterations. Differential expression was performed between normal and tumoral cells, between clone 1 and clone 3 and between clone 2 and aggregated clone 1 and 3. Differential expression was performed in the same way as in the breast xenograft example. We note that this two-steps clustering leads to results that are consistent with the solution obtained with $k = 4$ if we do not remove tumour cells from the input.

#### Hematopoietic precursors data analysis

Data for 4 healthy donors and 5 patients with bone marrow failure were available from (Zhao et al. 2017). We first performed standard quality check and normalisation using Seurat. In detail, we removed cells with a high percentage of counts coming from mitochondrial genes (cutoff > 15%) and with counts consistently lower or higher than the majority of the population (>6.8e+6 reads and <2.4e+7 reads).

Data was then normalised in CPM values (counts per million) and transformed in logarithmic scale through a $log(x + 1)$ transform. As not all the patients had CNA events we selected patient 1 with monosomy at 7 and an healthy individual as control (labelled with H4). As the dataset contained different cellular populations to remove the biases caused by marker genes highly expressed in just one population we first median filtered the gene expression counts for each chromosome using the “runmed” function in R, using a window dimension of 11.

Given the absence of a corresponding DNA-derived bulk CNA profile, and therefore lacking a segmentation associated, we had to resort to a custom segmentation, hopefully suitable to target chromosome-level aneuploidy events. We indeed segmented the genome at the level of whole chromosomes, and assumed a prior ploidy of 2 for each of them, which seems a reasonable baseline choice. After this we runned CONGAS using respectively 3.5 and 0.01 as scale and rate parameters for the library size factors prior; we had to change these values compared to other analyses as our defaults were optimised for 10x protocols. Clusters with less than 10 cells were discarded as outliers. Differential expression was performed with DESeq2 as for the other datasets.

As in the PDX example, we run copyKAT with default parameters. This time the dataset was composed of both normal and aneuploid cells, following the ideal experimental design for copyKAT. In this case the inference was extremely good, we clustered the cell tree and the aneuploid cluster was not splitted (Supplementary Figure S18).

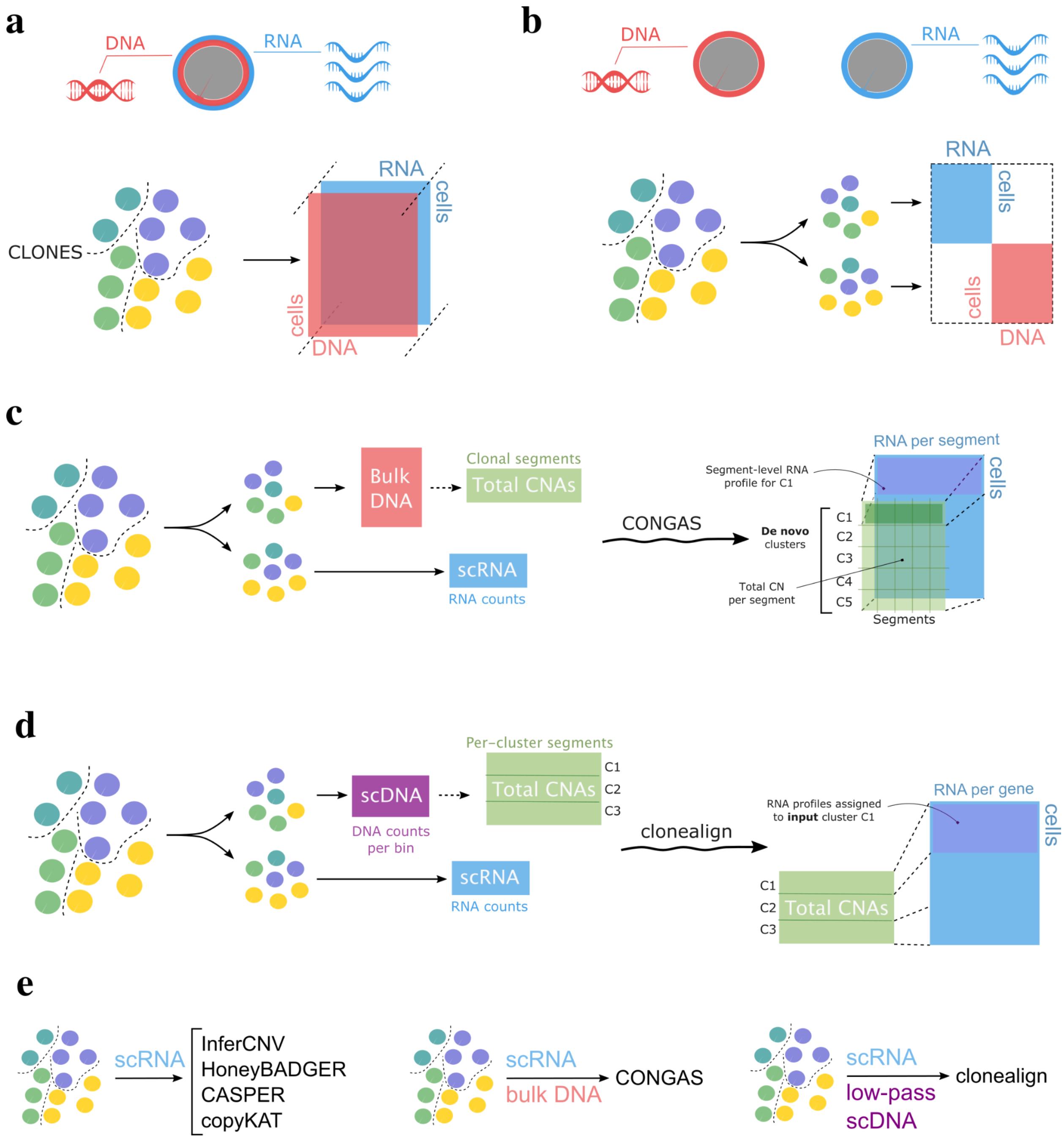


**Supplementary Figure S1. a.** Single-cell multi-omics from a population of heterogeneous cells, phylogenetically related into 4 clones (distinct colours). In this assay we simultaneously measure the DNA and RNA from the same cell, and use a tensor to describe the data. **b.** If we cannot implement a multi-omics assay, we can split the cells into groups and generate independent measures; these data cannot be directly overlapped. **c.** CONGAS takes in input clonal CNAs estimated from bulk DNA sequencing, which can be represented as a vector. Then it uses this vector to build a Bayesian prior and infer de novo CNAs in single-cells from scRNA data. The new clusters will have different copy number profiles (per segment), and will be used to aggregate cells with the same CNAs. **d.** Clonealign takes in input a matrix of CNAs per clusters, estimated from low-pass single-cell DNAseq (scDNA). Then, it assigns cells in scRNA-space to the clusters profiles given in input. This is the main difference with respect to CONGAS, which infers clusters de novo based on the input individual profile; the link function to predict RNA counts from segment ploidy is linear, and is the same for both methods. **e.** Methods to detect copy numbers from scRNA, alternative to CONGAS and clonealign. InferCNV, HoneyBADGER, CASPER and copyKAT detect CNAs from scRNA by segmenting RNA. These methods decouple CNA detection from clustering of cells, while CONGAS performs calling and clustering in a unique statistical model.

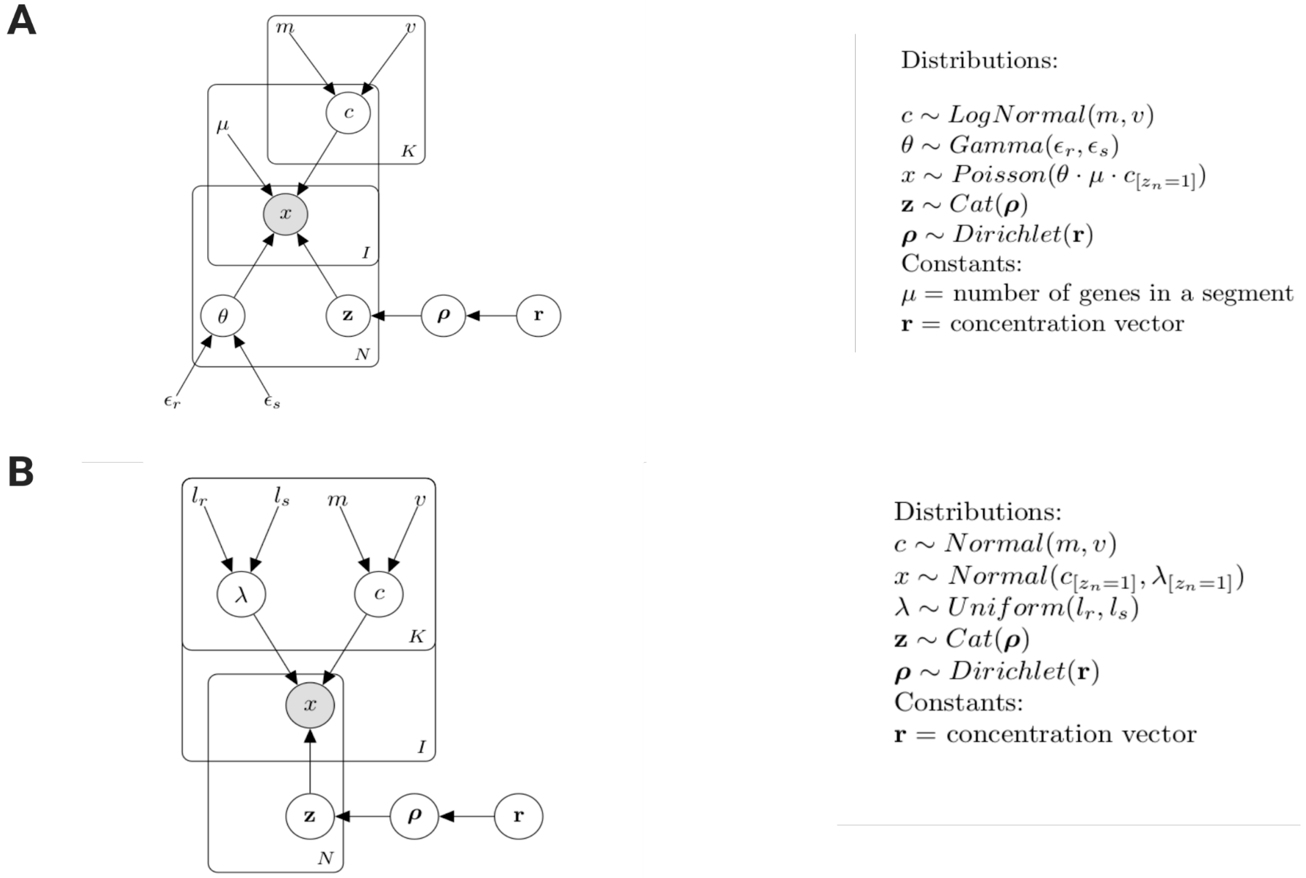


**Supplementary Figure S2.** CONGAS probabilistic graphical models in plate notation. **A.** CONGAS main model for counts as a finite mixture of Poissons; here $N$ indexes the number of cells, while $I > 0$ and $K > 0$ represent respectively the total number of segments and clusters. Note that latent variables **z** are vectors of dimension $K$ to represent clustering assignments (per cell). **B.** CONGAS alternative model as a finite mixture of independent Gaussian distribution, which can process continuous inputs. We assume in this case that the data is not only discretized, but has already been normalised for library size and other confounding variables.

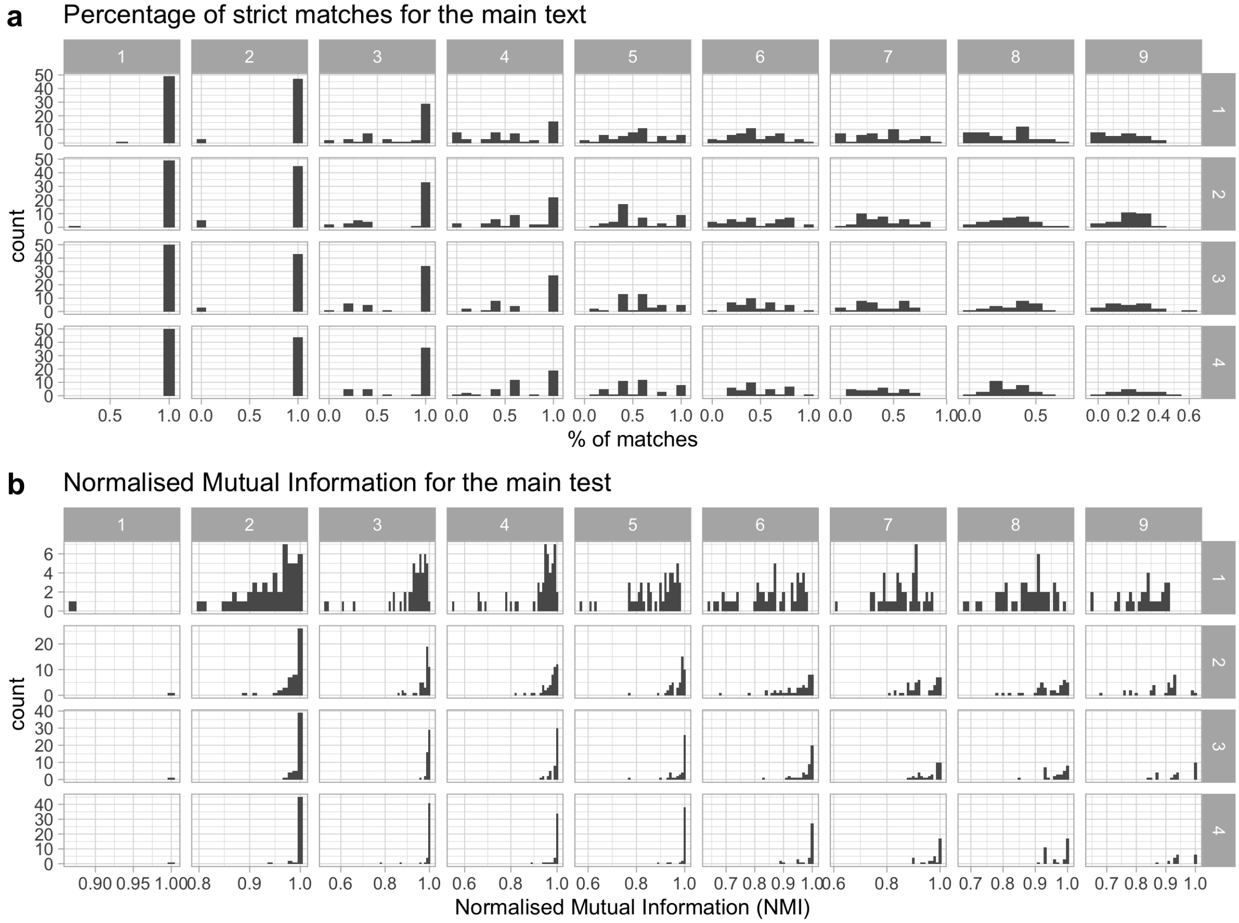


**Supplementary Figure S3. a,b.** Performance for the main synthetic test, where we scan different numbers of clusters assembled on trees with increasing evolutionary distance. We show the percentage of cluster labels that match in a simulation. We do this by sorting clusters by size, assigning them labels, and then assigning each cell to a cluster according to the model. We then check position by position if the clusters match - this is a strong penalty. In bottom we show NMI, the mutual information normalised in [0,1].

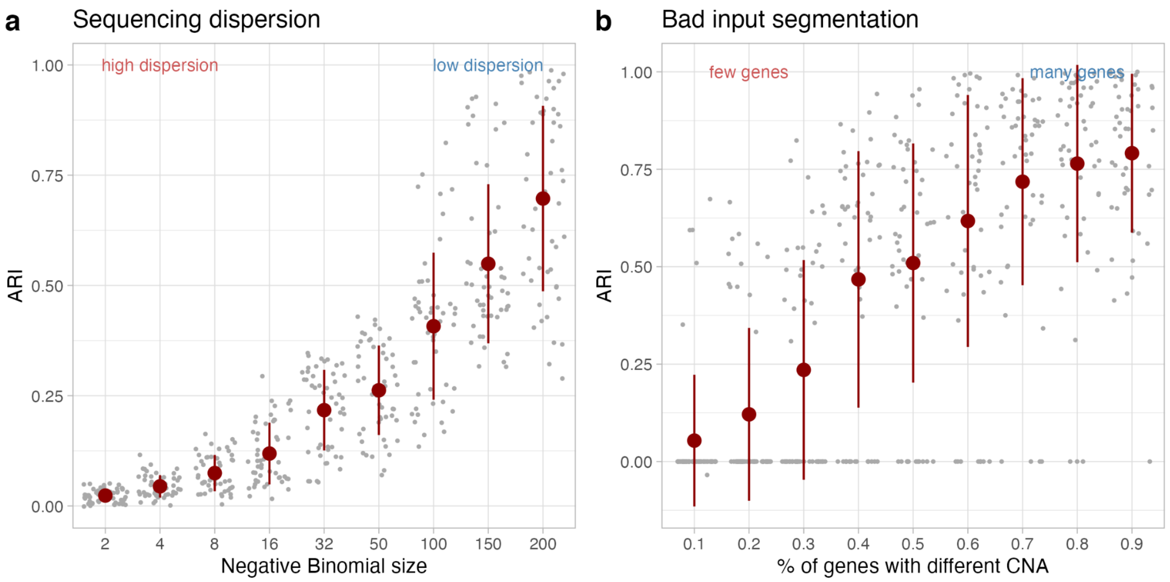


**Supplementary Figure S4.** **a.** ARI from synthetic tests simulating sequencing overdispersion after changing the Poisson model in CONGAS with a Negative Binomial and variable dispersions. For each test a clonal architecture with 2 clones. is simulated. **b.** ARI from synthetic tests simulating input misleading segmentations, meaning that only a subset of genes that map to a segment is affected by the CNA, i.e., the input breakpoints from bulk do not match real breakpoints in single-cell data. For each test a clonal architecture with 2 clones and a fixed number of 2 segments is simulated.

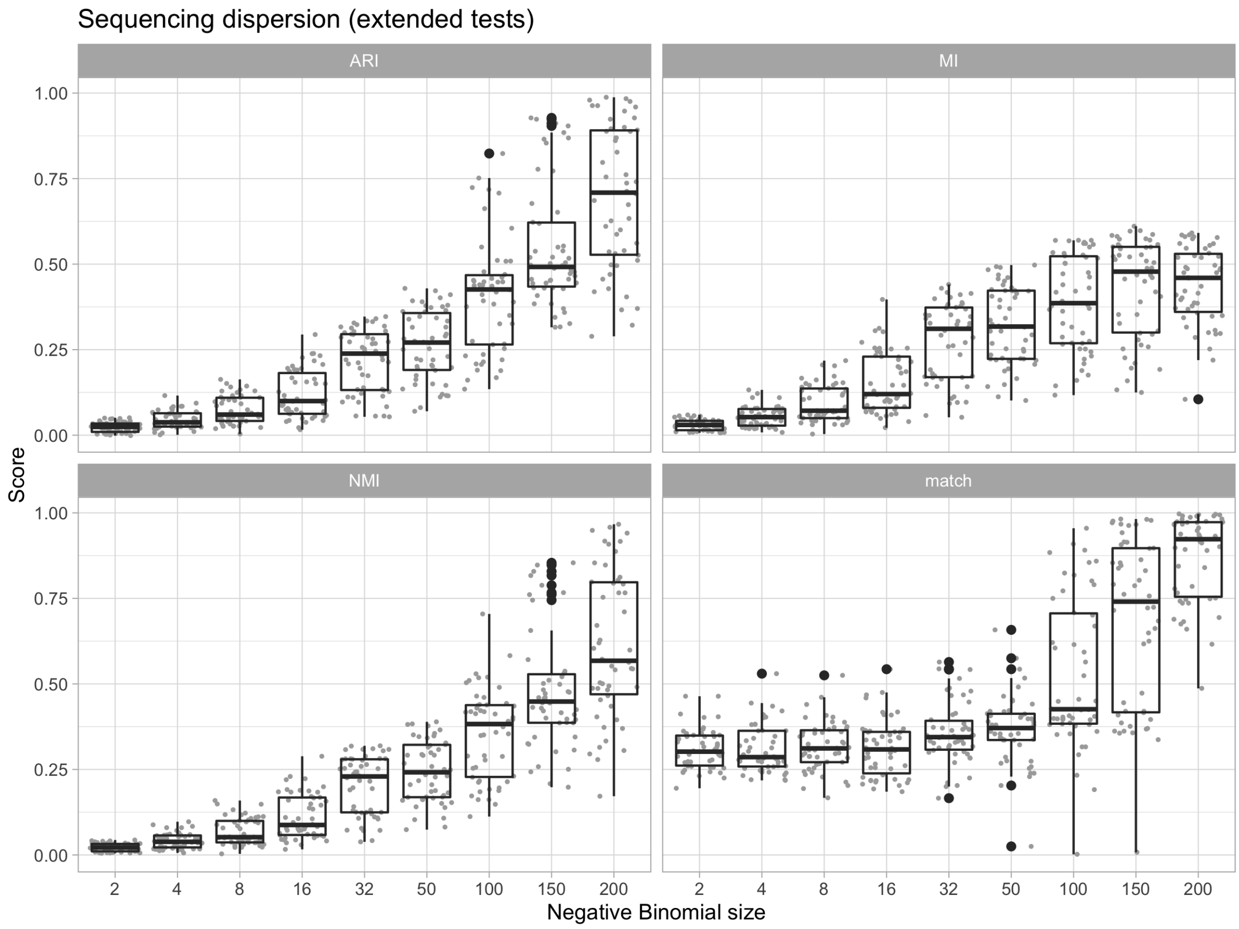


**Supplementary Figure S5.** Performance for synthetic tests with sequencing overdispersion. In this case we use different parameters of a Negative Binomial distribution to generate read counts (Main Text). The performance shows a clear trend; here the scores are computed between simulated and inferred clustering labels. ARI is the adjusted rand index, MI and NMI the mutual information and its normalised extension. Score match reports the proportion of cells with the exact same cluster label (simulated versus inferred).

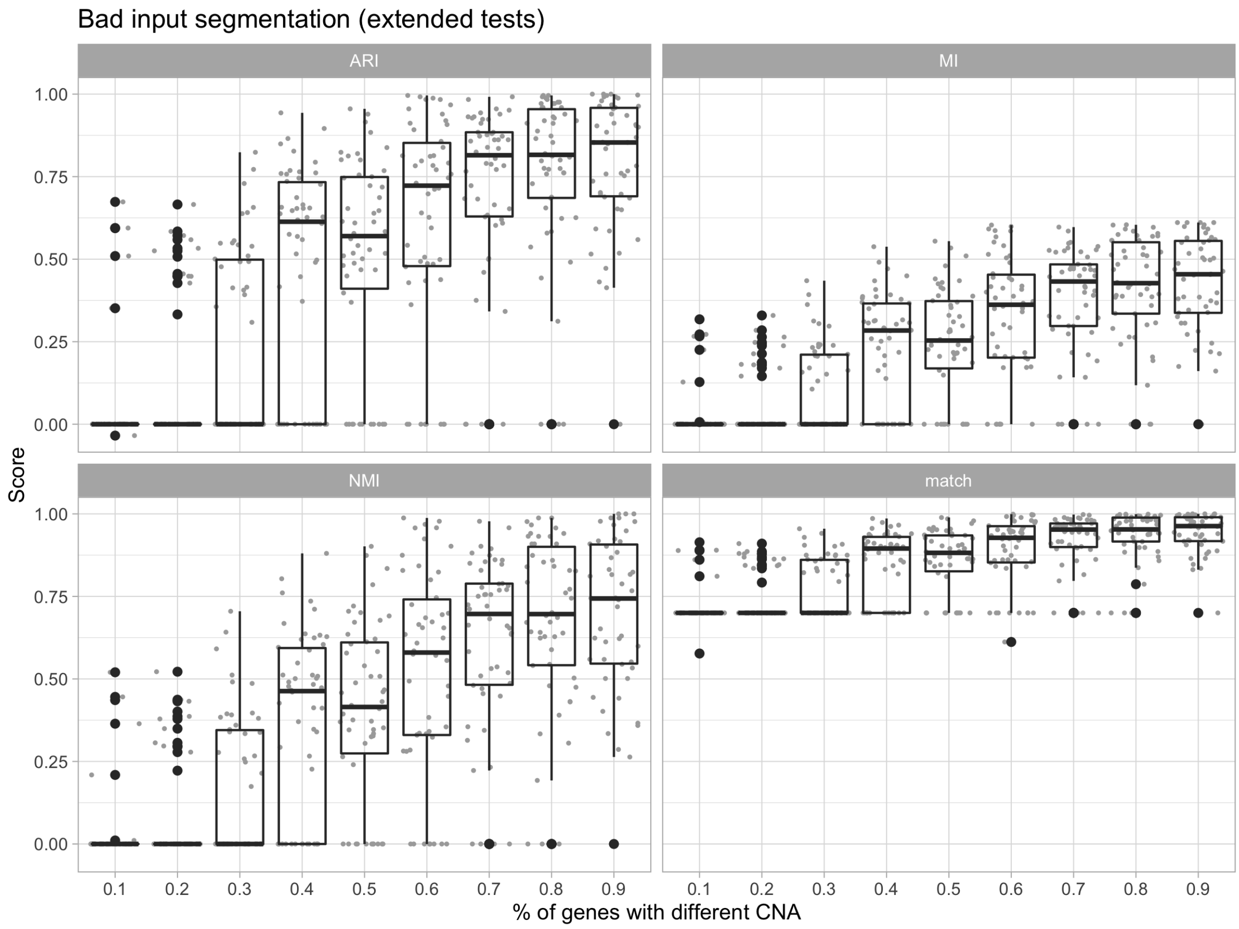


**Supplementary Figure S6.** Performance for synthetic tests with subclones that have CNA segments where only a portion of the mapped genes obeys the linear DNA/RNA relation. In practice, this tests for the presence of subclonal CNAs that involve segments smaller than the ones given in input to CONGAS. In this case we use different proportions of genes, ranging from 10% to 90% (see also Main Text). The performance shows a clear trend; the same scores in Supplementary Figure S2 are reported.

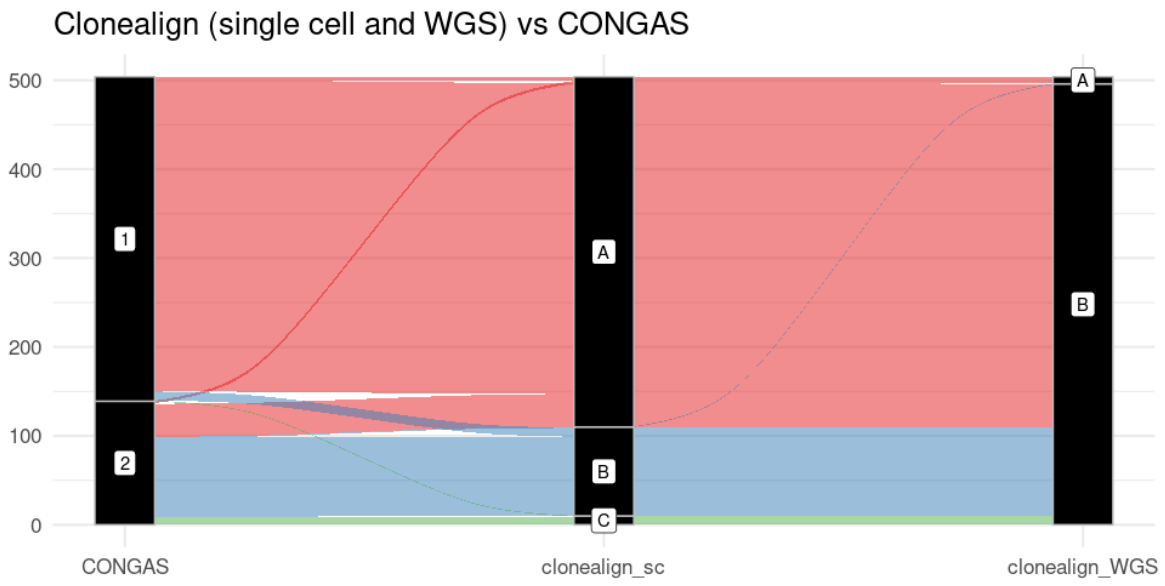


**Supplementary Figure S7.** Sankey plot showing the very strong agreement between the clusters identified by CONGAS and the clusters A and B identified by scDNA-seq and assigned with clonealign in the original paper (clonealign_sc in figure). The assignments in the left part (clonealign_WGS) are obtained by resolving the subclonal structure directly from the bulkDNA WGS of the primate tissue and using ReMixT, a tool capable of finding subclonal CNAs from bulk DNA. As you can see in this case clonealign is not able to distinguish the two clones and assigns almost all cells to clone B.

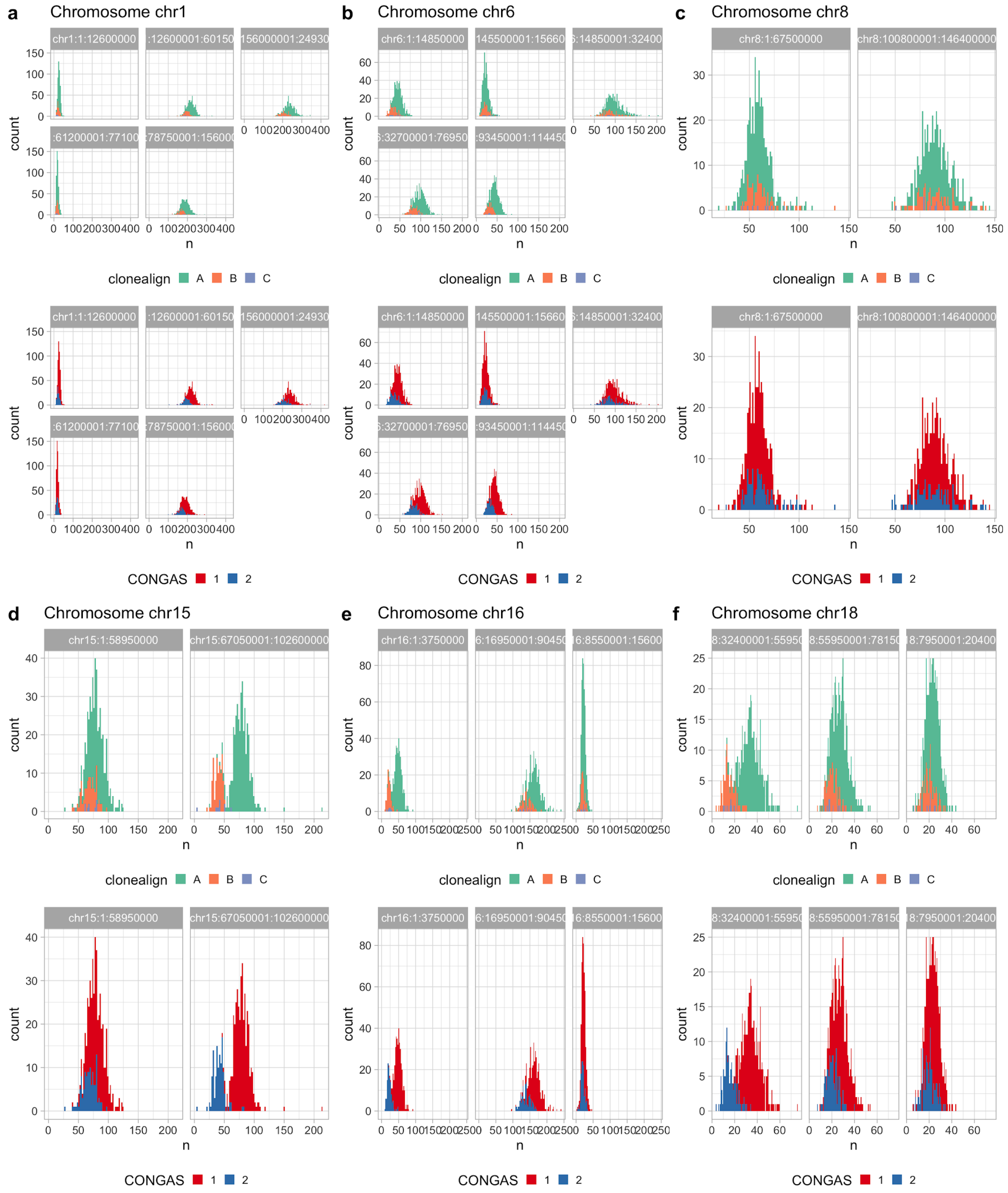


**Supplementary Figure S8. a-e.** For every segment in a subset of the input chromosomes we plot the data coloured accordingly to the clustering assignments obtained by clonealign (top), and CONGAS (bottom). Clone C from clonealign is very difficult to identify; both other clusters match perfectly

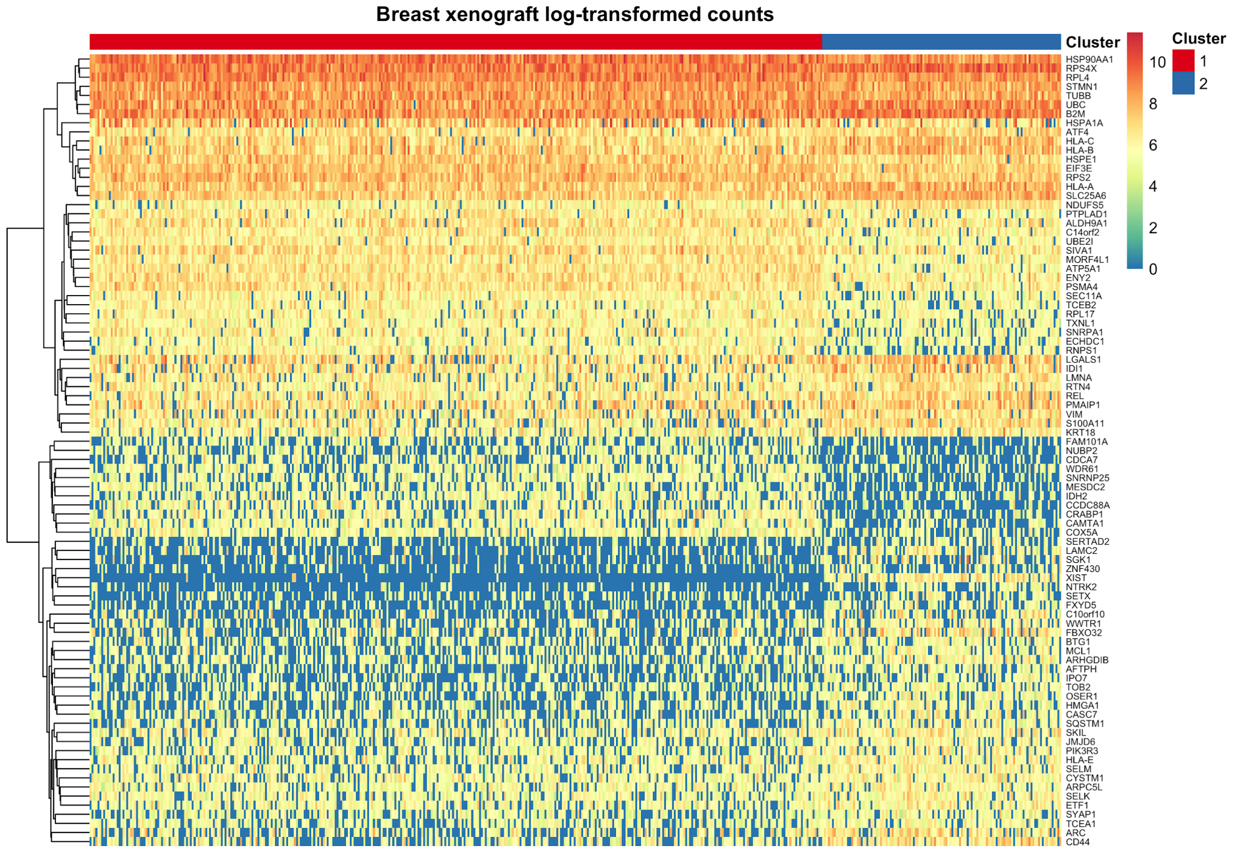


**Supplementary Figure S9.** Heatmap for the input raw counts of the breast cancer xenograft discussed in the Main Text. Each column is a cell, each row one of 212 differentially expressed genes; count values are log transformed. Rows are clustered by hierarchical clustering.

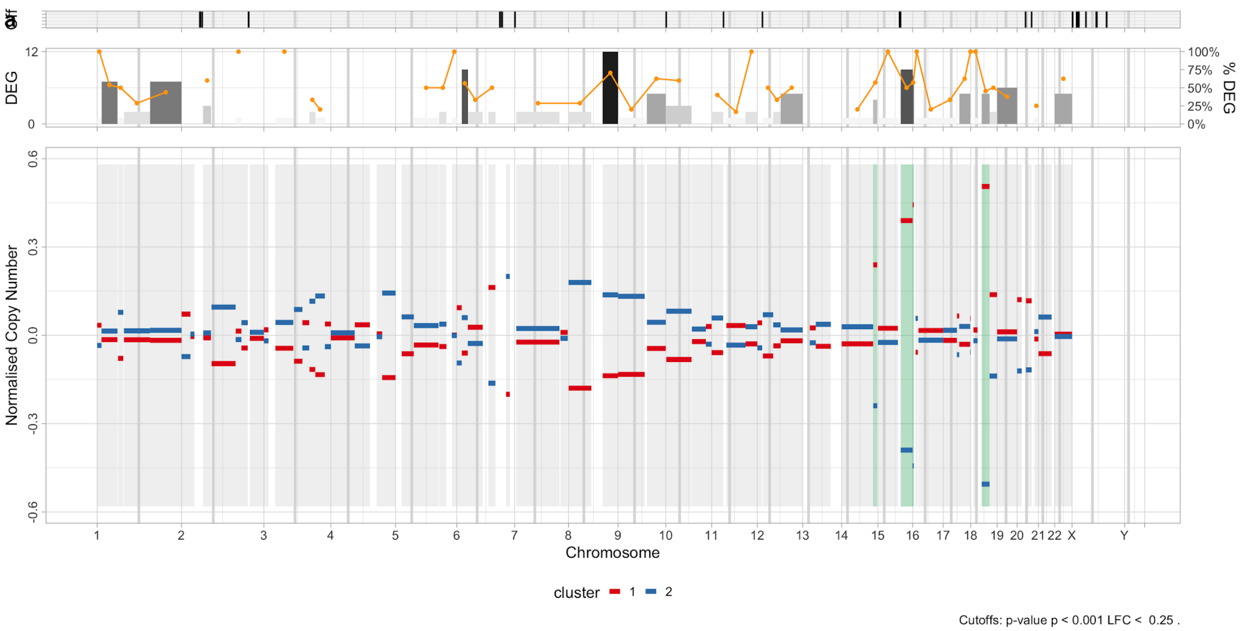


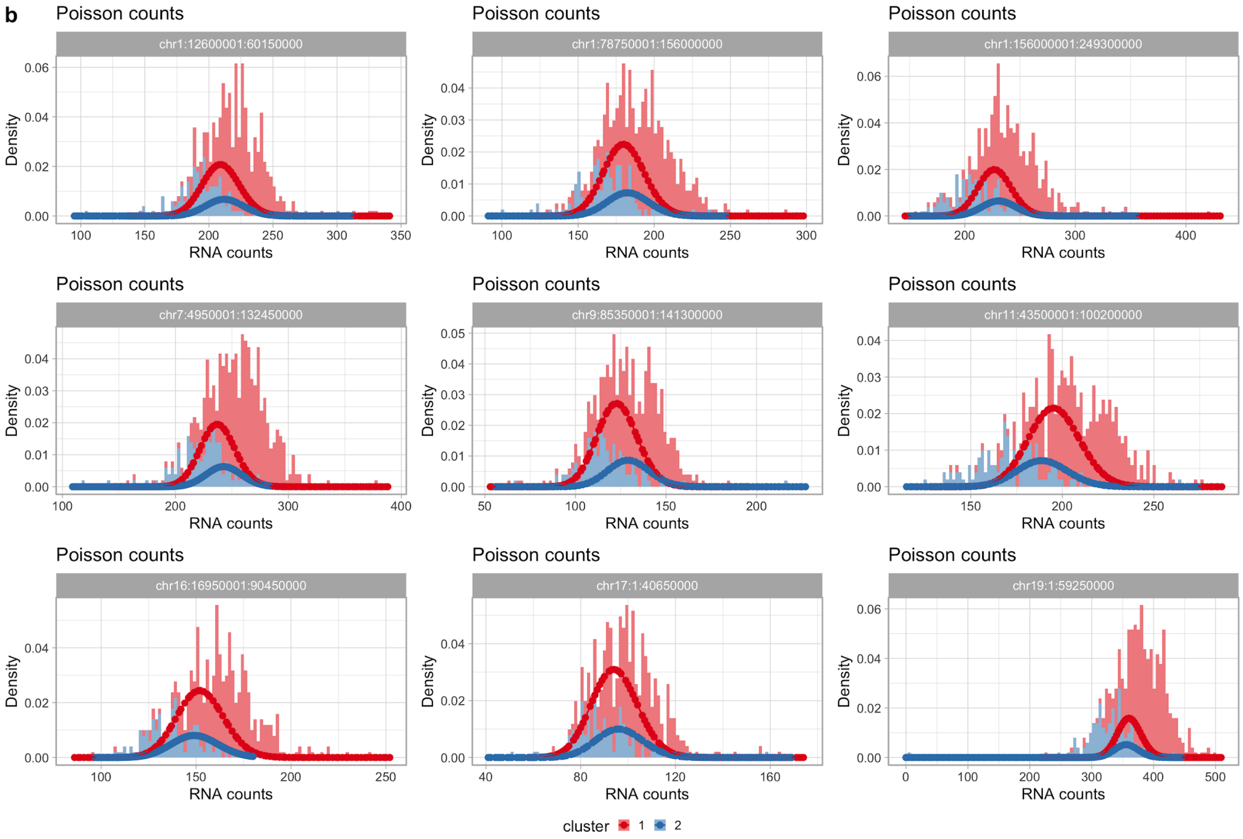


**Supplementary Figure S10. a.** Genome-wide DEGs for the breast cancer xenograft discussed in the Main Text. Above the genome we show the number and the percentage of DEGs per segment; the top marks are DEGs off the input CNA segments. **b.** Data density and CONGAS mixture for segments with >250 genes (the largest segments of this tumour).

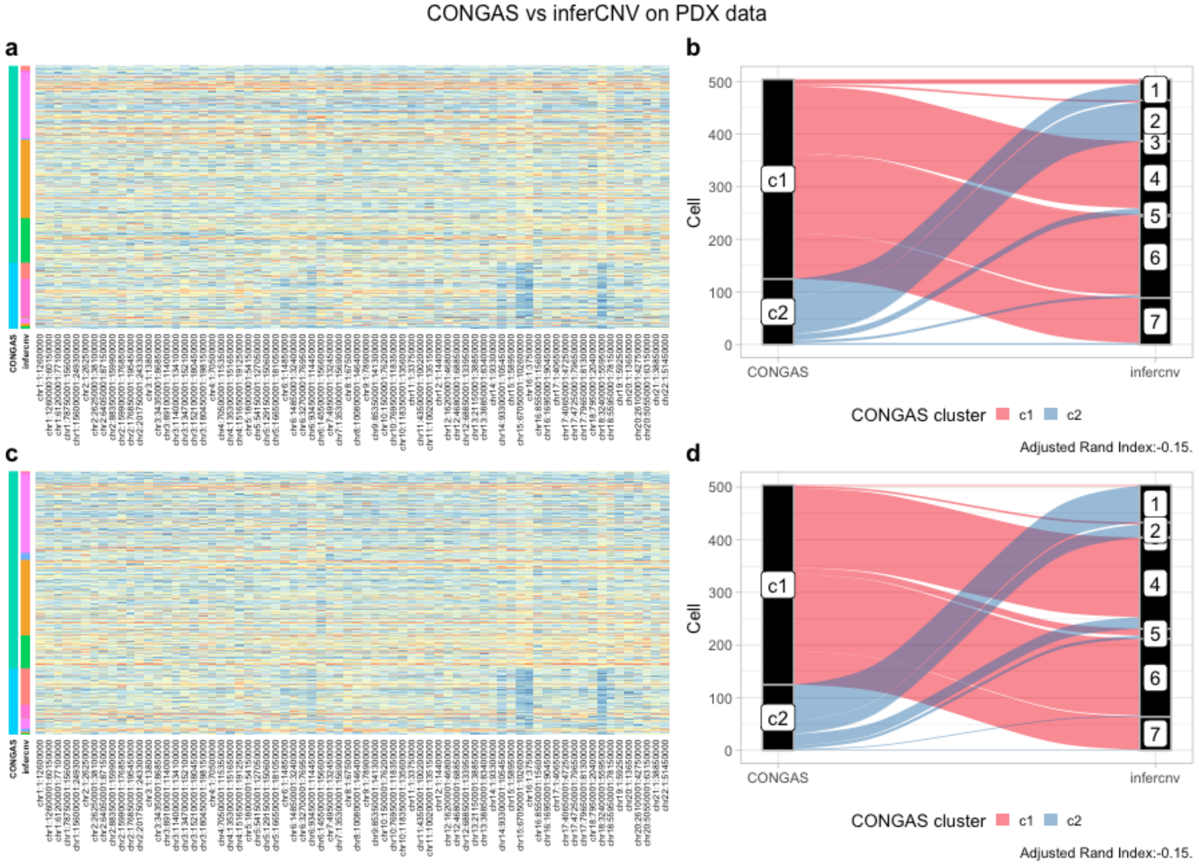


**Supplementary Figure S12. a.** Heatmap of segment counts (z-score transformed) ordered by inferCNV clustering assignments for the PDX example dataset. In this case inferCNV was runned without a reference **b.** Sankey plot for clustering assignments of CONGAS and inferCNV (no reference), the c2 cluster of CONGAS is actually splitted in two smaller clusters, while the other clone is splitted in 5 parts. Confirming the tendency of overclustering of the random tree method implemented in inferCNV **c.** Heatmap of segment counts (z-score transformed) ordered by inferCNV clustering assignments for the PDX example dataset runned with reference from GTEx (breast tissues only) **d.** Sankey plot for clustering assignments of CONGAS and inferCNV (with reference), results are almost identical to the run without reference shown in panel (b)

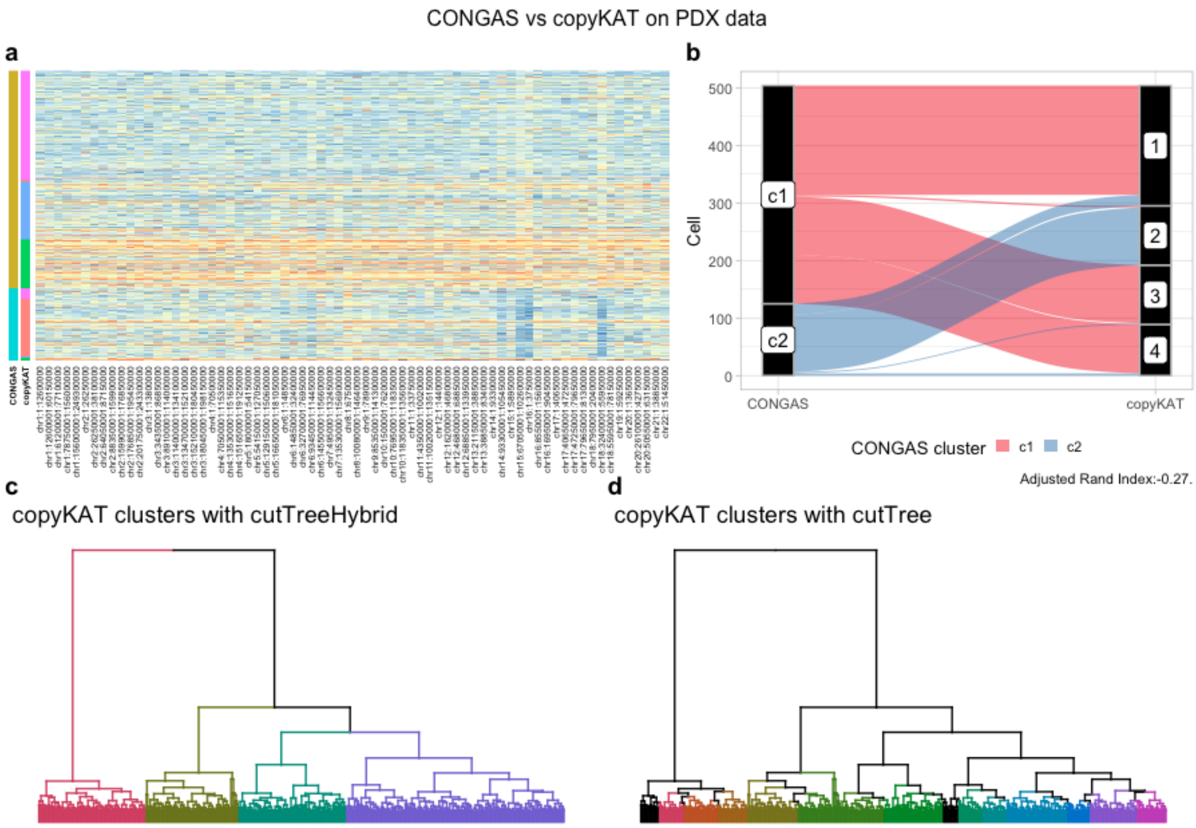


**Supplementary Figure S13. a.** Heatmap of segment counts (z-score transformed) ordered by copyKAT clustering assignments for the PDX example dataset. **b.** Sankey plot for clustering assignments of CONGAS and copyKAT. It can be clearly seen how cluster “c2” of CONGAS corresponds almost entirely to copyKAT “cluster2”, while the bigger cluster “c1” is divided into 3 smaller clusters by copyKAT with cutTreeHybrid. **c.** Automatic tree cutting using the hybrid algorithm of dynamicTreeCut (default) **d.** Automatic tree cutting using the “tree” algorithm of dynamicTreeCut.

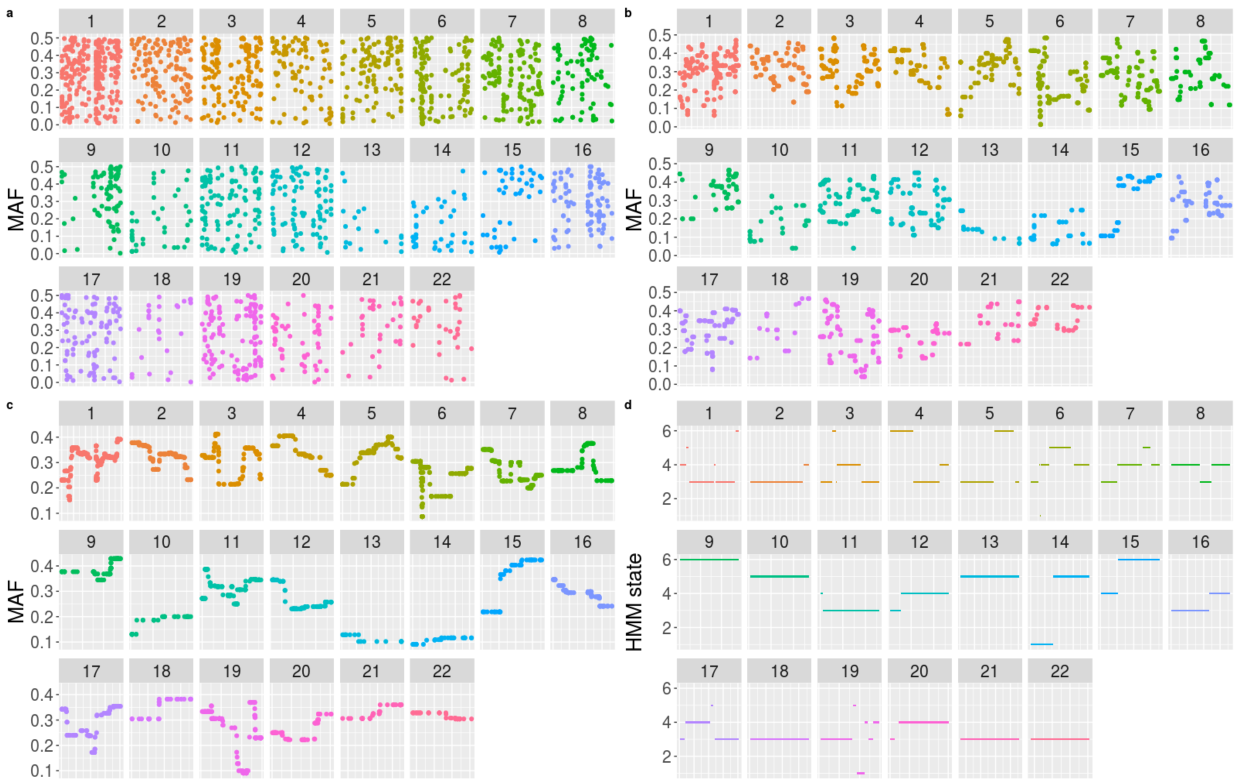


**Supplementary Figure S14.** Results of the first run for our HMM semgenter **a-c.** Plots of the raw MAF data after median filtering over chromosomes. The three plots show increasing width for the median window, in order from the upper left 1 (no filtering at all), 5, 21. These plots also highlight the intrinsic noise in the data, which makes it hard to call confidently the CNV regions **d.** Plot of the HMM states after inference provide breakpoints and segmentation values.

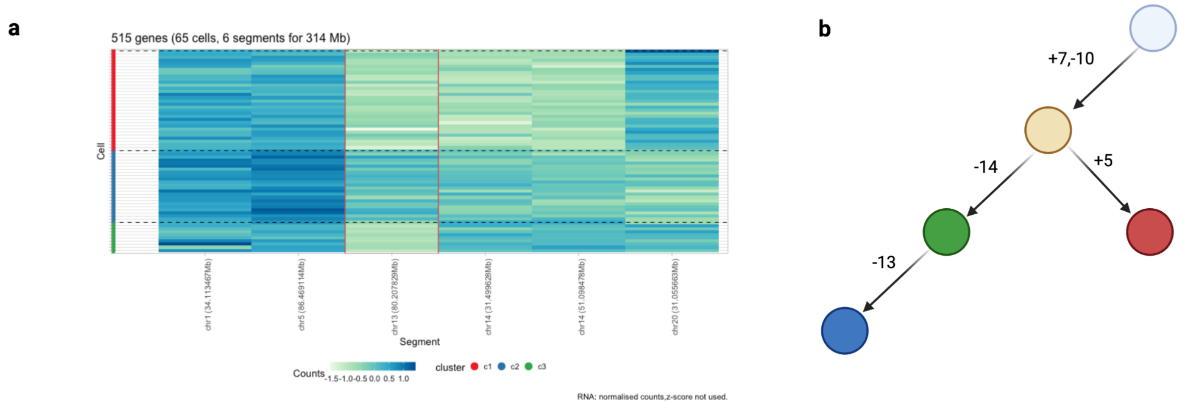


**Supplementary Figure S14. a.** Z-score normalised segment values for CONGAS run on GBM data after normal cells have been removed, the clustering assignment is the same as the panel c-d in Figure 3 **b.** Maximum parsimony tree for the GBM dataset**.** The +7,-10 alteration can be found in the comparison against normal cell (the first run of clustering) and are common to all the cell of the tumours, then there is a branching where we found a clone with a -14 (c3), and another one with a +5 (c2). The last leaf on the right (clone c1) derives from c3 after a deletion on chromosome 13.

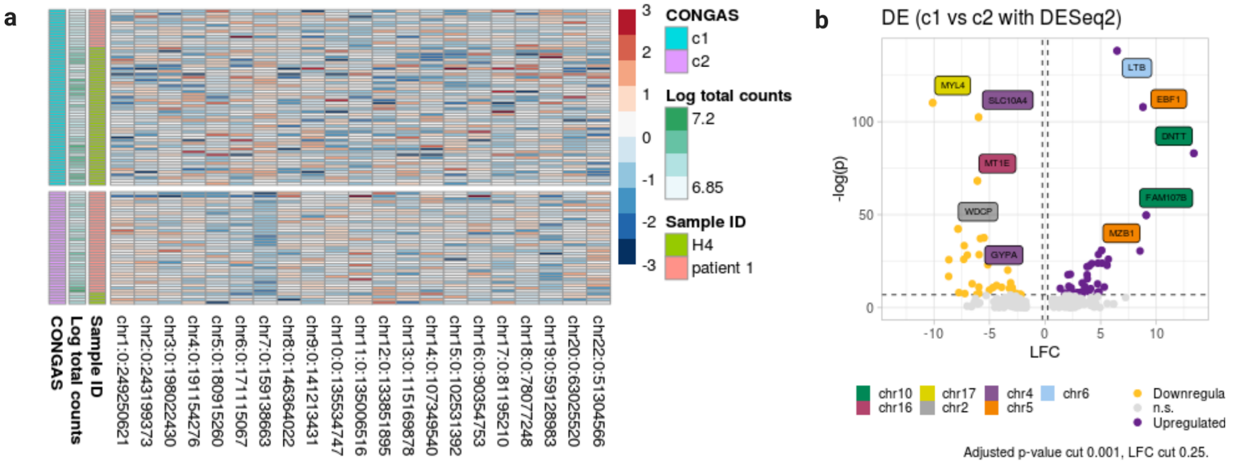


**Supplementary Figure S15. a.** Analysis of bone marrow Smart-Seq samples for $n = 100$ cells and 2 patients, one healthy (H4) and one with bone marrow failure (patient 1). Since this dataset does not have a precalculated input CNA segmentation for CONGAS, we have aggregated gene counts at the level of whole chromosomes. This panel shows how CONGAS distinguishes between the healthy population and the one with the disease, due to a deletion in chromosome 7 in some cells of patient 1. It can also be noticed how clustering assignments are not correlated with the total number of counts, suggesting that CONGAS can correctly normalise for sequencing efficiency. **b.** Volcano plot showing the gene differentially expressed between the two clusters; note that the highlighted genes map off from chromosome 7.

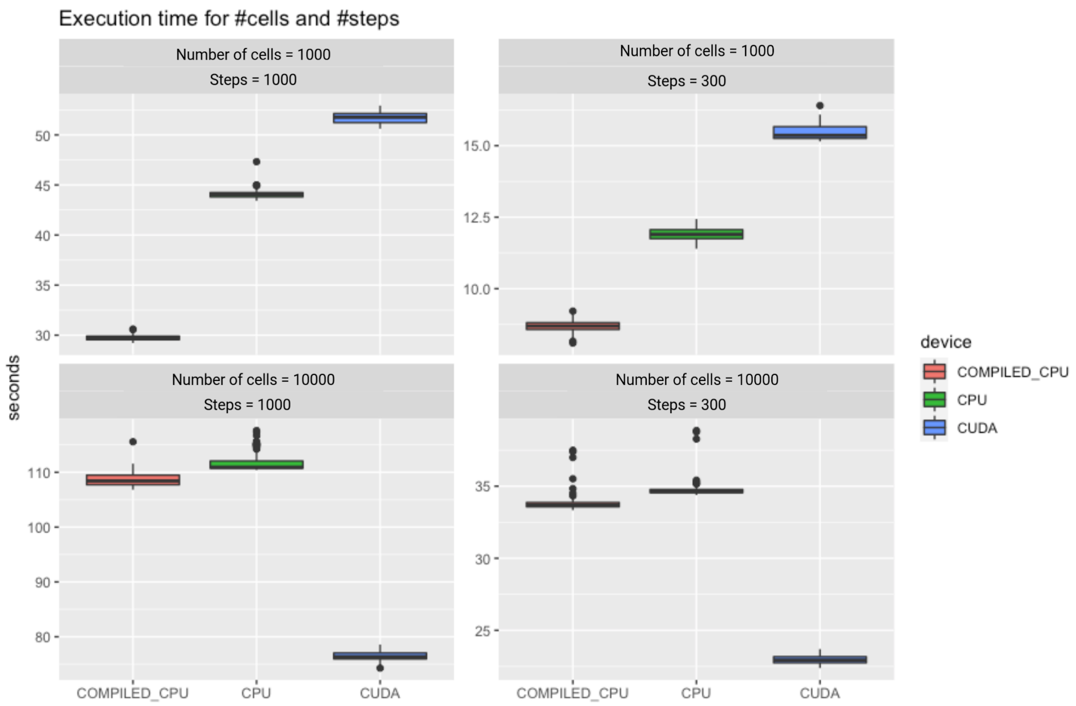


**Supplementary Figure S15.** Execution time for CONGAS runned on two simulated dataset, one with 1000 cells and the other with 10000 cells. For each dataset we timed 100 executions for respectively 300 and 1000 gradient update steps in 3 different settings: standard python interpreter, JIT compiler and GPU/CUDA. From the plot it is clear how for small cell numbers the CPU is faster, but as the number of cells starts to increase the GPU can give an effective speed up to the calculation. All calculations were performed on a machine with 2 Intel Xeon vCPU @2.2GHz, 13 GB of RAM and an NVIDIA Tesla T4.

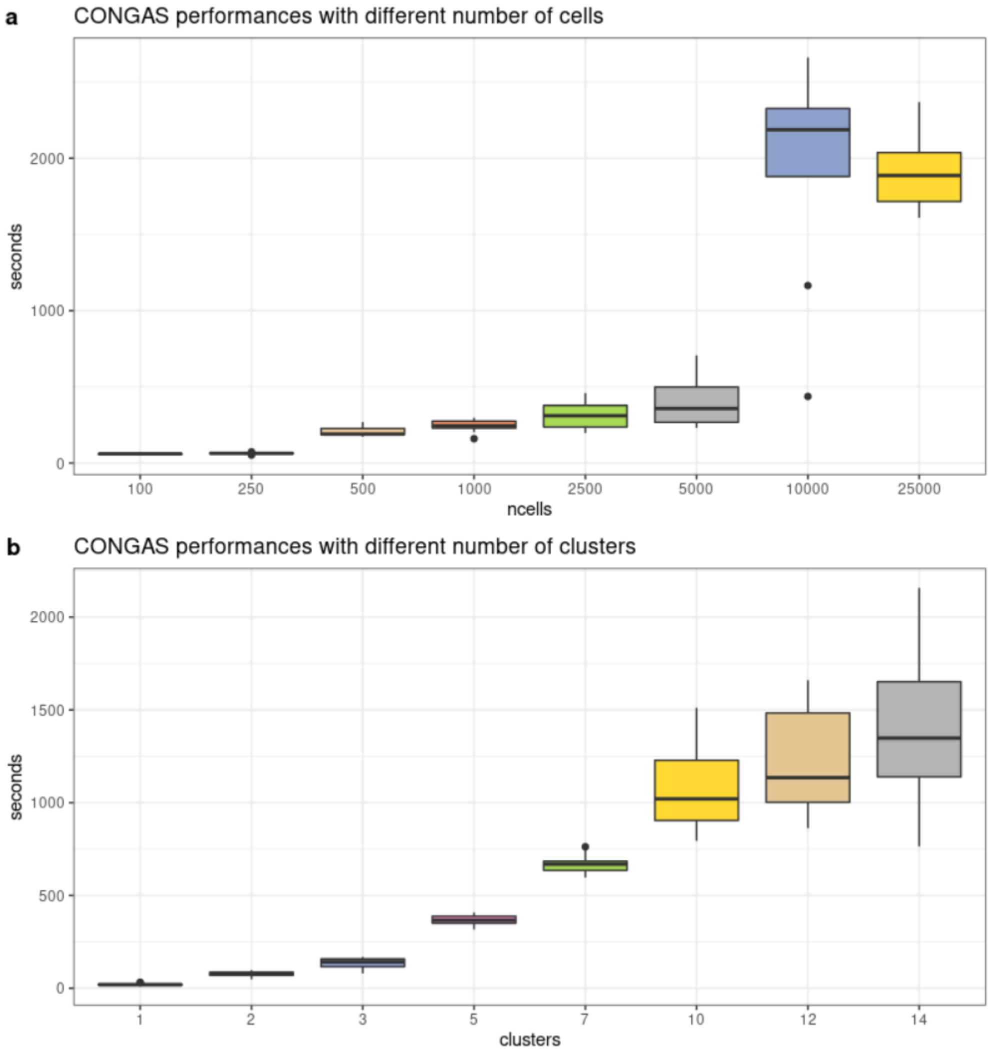


**Supplementary Figure S16. a.** Execution time for CONGAS runned on a simulated dataset with a number of cells ranging from 100 to 25000. Each box corresponds to 10 repetitions. Timings are for just a single run with K=5. Here the trend is discontinuous with execution times almost equal till a discrete jump, this is probably due to the memory usage routine of PyTorch **b.** Execution time for CONGAS runned on a simulated dataset with K ranging from 1 to 14. Each box corresponds to 10 repetitions. Timings are for a number of cells equal to 500.

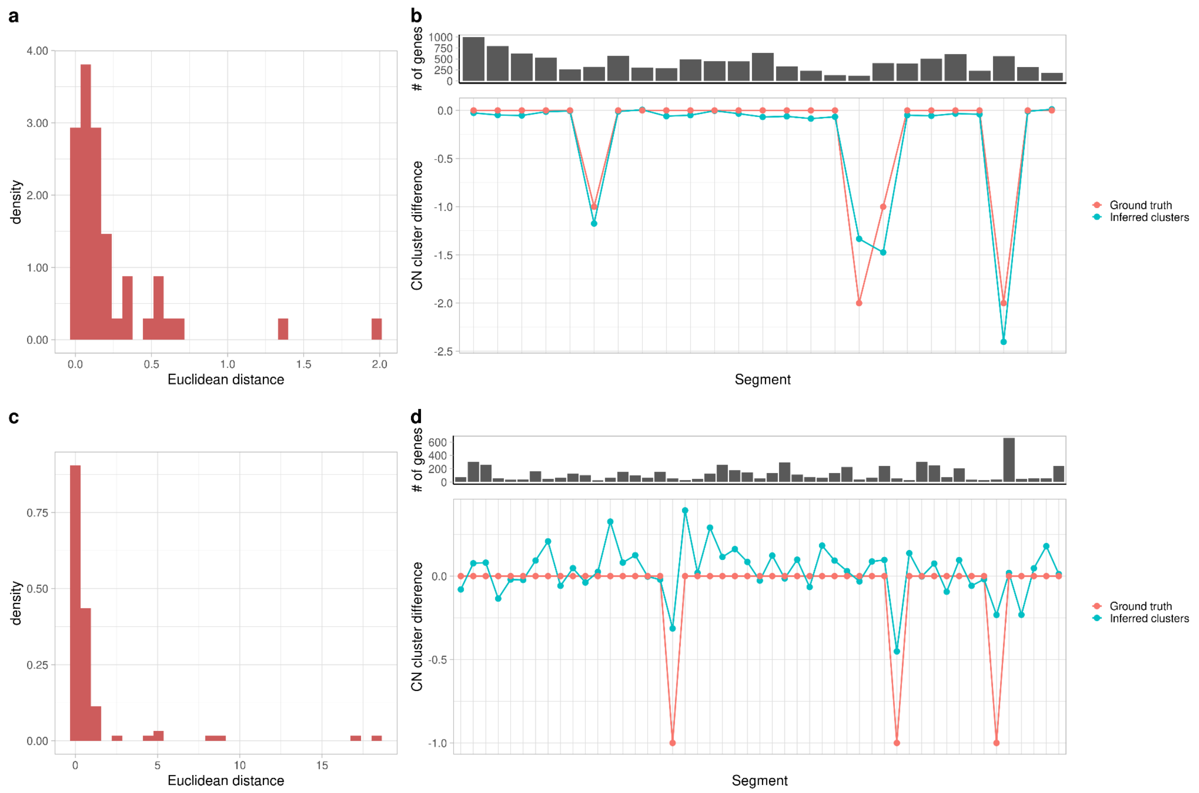


**Supplementary Figure S17. a-c.** euclidean distance between the inferred copy number profiles and the corresponding ground truth profile for each cluster in a simulated instance with two clusters (a) and in the triple-negative breast xenograft dataset (c). **b-d.** difference between the profiles of the two clusters, both for the ground truth and the inferred clones. Results are displayed for a simulated instance with two clusters (b) and the triple-negative breast xenograft dataset (d).

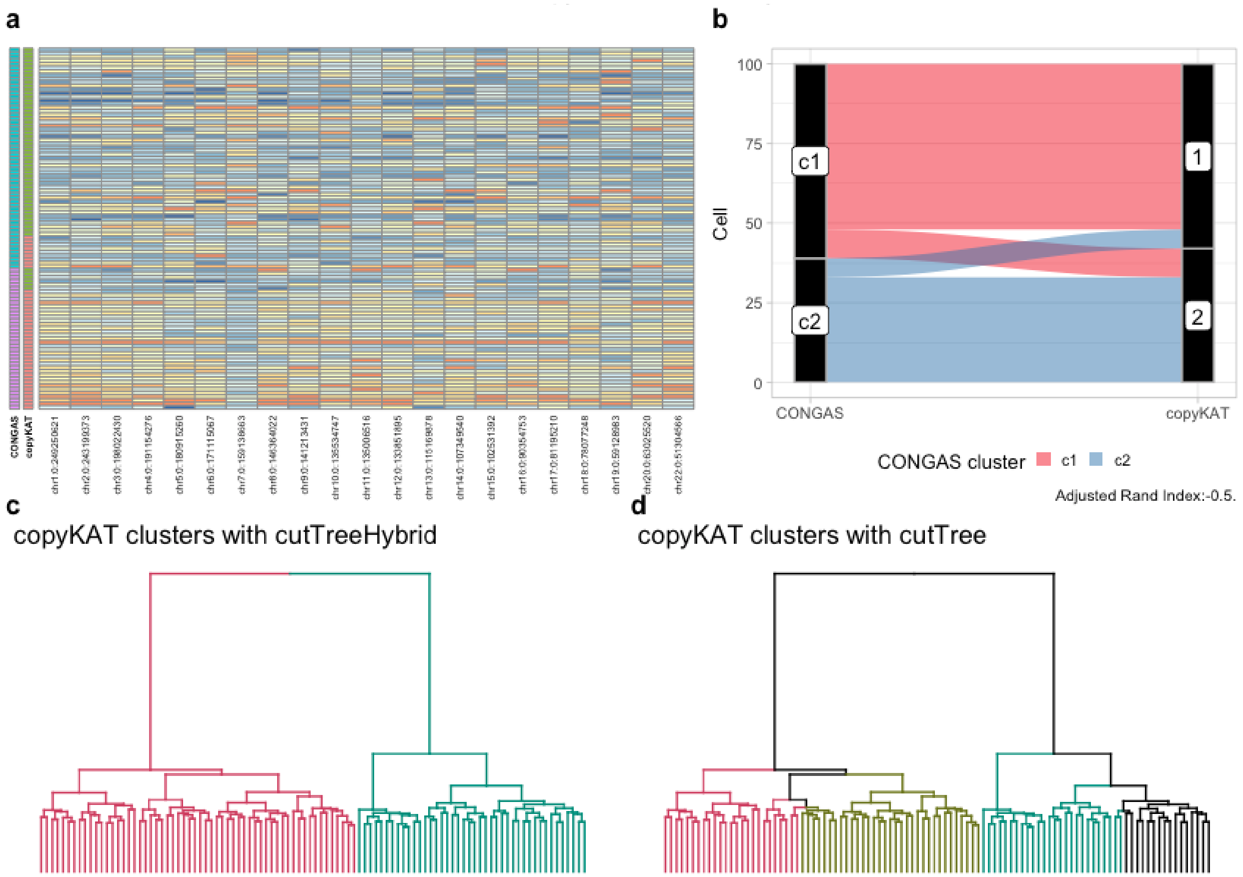


**Supplementary Figure S18. a.** Heatmap of segment counts (z-score transformed) ordered by copyKAT clustering assignments for the Monosomy 7 dataset. **b.** Using the Sankey plot for clustering assignments of CONGAS and copyKAT, we can clearly see how there is an extremely good consensus between the two algorithms. **c.** Automatic tree cutting using the hybrid algorithm of dynamicTreeCut, which also takes into account the distance matrix, this is the default used in the panels a-b and across the paper **d.** Automatic tree cutting using the “tree” algorithm of dynamicTreeCut, which uses just the topological information.
